# Encoding of the colorectal cancer metabolic program through MICU2

**DOI:** 10.1101/2023.10.31.564893

**Authors:** Alison Robert, David Crottès, Jérôme Bourgeais, Naig Gueguen, Arnaud Chevrollier, Jean-François Dumas, Stéphane Servais, Isabelle Domingo, Stéphanie Chadet, Julien Sobilo, Olivier Hérault, Thierry Lecomte, Christophe Vandier, William Raoul, Maxime Guéguinou

## Abstract

The mitochondrial Ca^2+^ uniporter (MCU) plays crucial role in intramitochondrial Ca^2+^ uptake, allowing Ca^2+^-dependent activation of oxidative metabolism. In recent decades, the role of MCU pore-forming proteins has been highlighted in cancer. However, the contribution of MCU-associated regulatory proteins mitochondrial calcium uptake 1 and 2 (MICU1 and MICU2) to pathophysiological conditions has been poorly investigated. Here, we describe the role of MICU2 in cell proliferation and migration using *in vitro* and *in vivo* models of colorectal cancer (CRC). Transcriptomic analysis demonstrated an increase in MICU2 expression and the MICU2/MICU1 ratio in advanced CRC and CRC-derived metastases. We report that expression of MICU2 is necessary for mitochondrial Ca^2+^ uptake and quality of the mitochondrial network. Our data reveal the interplay between MICU2 and MICU1 in the metabolic flexibility between anaerobic glycolysis and OXPHOS. Overall, our study sheds light on the potential role of the MICUs in diseases associated with metabolic reprogramming.

**Highlights:**

- MICU2 plays a pivotal role in the balance between anaerobic glycolysis and oxidative phosphorylation
- MICU2 stabilizes the mitochondrial network and endoplasmic reticulum–mitochondrial calcium flux
- MICU2 expression and the MICU2/MICU1 ratio control proliferation and metastasis formation in colorectal cancer

**Graphical Abstract:** 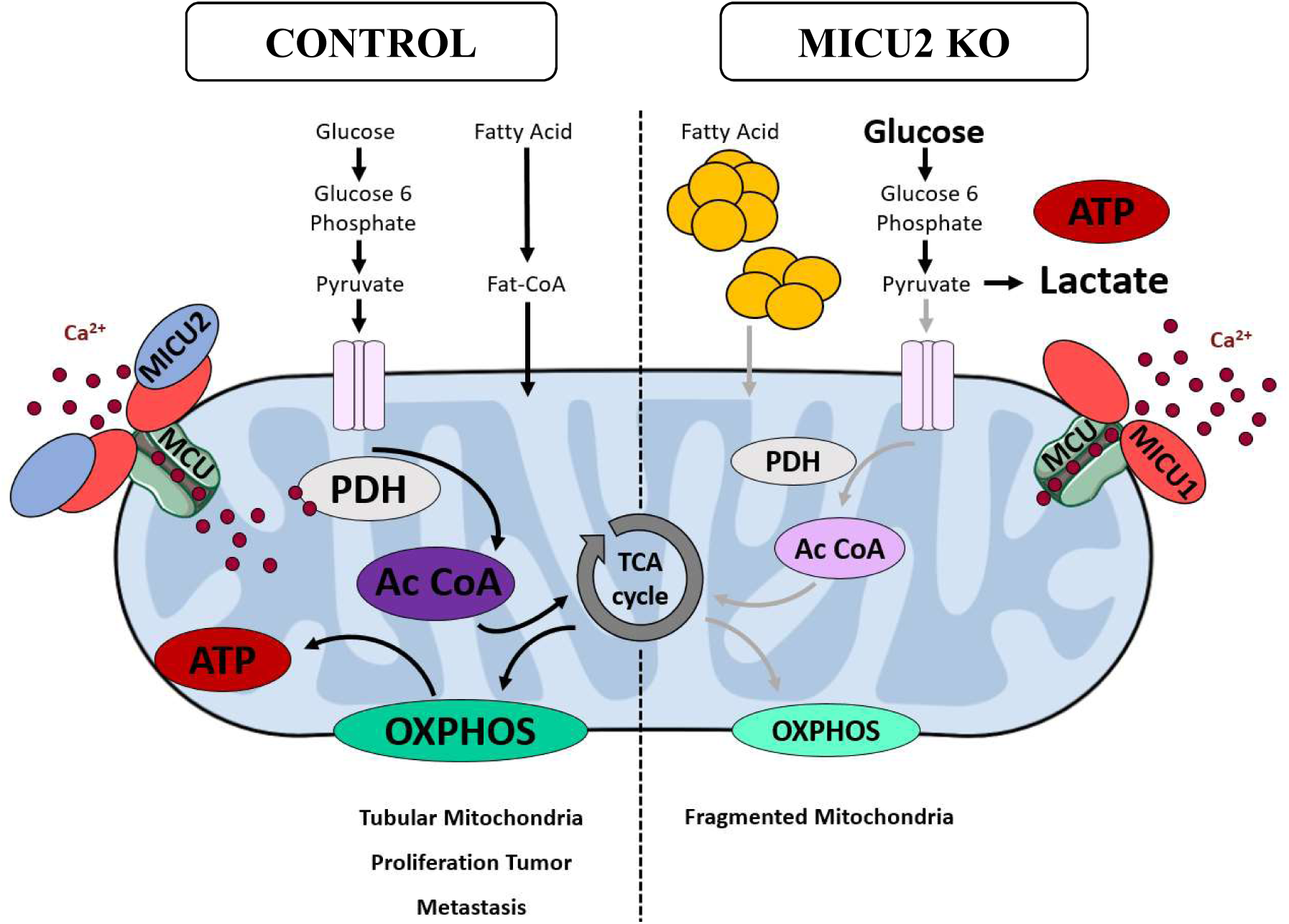

## Introduction

Intracellular calcium (Ca^2+^), which is a ubiquitous second messenger, regulates a multitude and various cellular processes. Deregulation of Ca^2+^ homeostasis and Ca^2+^ dependent signaling pathways have been described to be associated with each of the “cancer hallmarks”^1,2^. Mitochondria are a central player in maintaining cellular Ca^2+^ homeostasis by uptaking Ca^2+^ from cytoplasm.

Fine regulation of Ca^2+^ fluxes in mitochondria is required to support functions linked to metabolism (ATP production), redox homeostasis and proliferation/apoptosis balance. Mounting evidence indicates that metabolic transformation of tumor cell is the main limitation of cancer treatment, which is highly related to the resistance to therapeutic drugs^3^. Due to its role in energetic metabolism by activating pyruvate dehydrogenase, isocitrate dehydrogenase, and α-ketoglutarate dehydrogenase, or respiratory complex activity, mitochondrial Ca^2+^ (mitoCa^2+^) plays a pivotal role in cancer progression^4–7^.

To enter in the mitochondrial matrix, Ca^2+^ first passes through the outer membrane of the mitochondrion via the voltage-dependent anion-selective channel (VDAC), then through the inner mitochondrial membrane (IMM) via the mitochondrial calcium uniporter (MCU), a highly selective Ca^2+^ channel. MCU forms a large multi-molecular complex composed of different proteins. This channel, alongside the MCU regulator EMRE, are located in the IMM. Proteins regulating mitoCa^2+^ uptake such as mitochondrial calcium uptake 1 (MICU1), MICU2, and MICU3 reside in the intermembrane space.^8,9^ The regulation of MCU activity by MICU1 and MICU2 involves a gating mechanism. MCU gating and the role of MICU subunits in potentiation are dependent on the Ca^2+^ content and the MCU current. Recently, Garg et al.^10^ demonstrated that elevation of cytosolic [Ca^2+^] increases Ca^2+^ binding to the EF hands of MICU subunits. MICUs thereby increase the open probability of MCU, enhancing its activity. However, the contribution of each of these proteins in carcinogenesis is still poorly understood.

Colorectal cancer (CRC) is the third most common malignancy worldwide and one of the deadliest cancers. CRC involves a multi-step process that includes three distinct phases: initiation, progression, and metastasis. Researchers recently showed that MCU expression is increased and associated with poor prognosis in patients with CRC.^11^ Upregulation of MCU increases mitoCa^2+^ uptake to promote mitochondrial biogenesis and to facilitate CRC cell growth *in vitro* and *in vivo*.^11^

The role of MICU2 in the regulation of MCU activity depends on its binding to Ca^2+^ and is mediated by its binding to MICU1.^12^ MICU2 is a paralog of MICU1—it shares 42% amino acid identity with MICU1—and contains an amino-terminal mitochondrial targeting sequence and a cytosolic C terminus with two EF hands.^13^ Interestingly, the expression of MICU2 chiefly depends on MICU1, but the reciprocity has not been proved.^14,15^ So far, the contribution of MICU2 to the regulation of mitoCa^2+^ signaling, bioenergetics, metabolic reprogramming, mitochondrial dynamics, and CRC development have not been addressed.

Here, we observed that MICU2 expression and the MICU2/MICU1 ratio in patients with CRC are tightly correlated to CRC aggressiveness and stage. For the first time, we demonstrated that MICU2 regulates proliferation and migration of CRC cells *in vitro* and *in vivo*. We demonstrated the role of MICU2 on mitoCa^2+^ uptake and its contribution as a crucial element of mitochondrial function, regulating fusion-fission processes, ensuring the proper use of pyruvate by mitochondria and oxidation of fatty acid, and allowing the proper functioning of respiratory chain complexes. Overall, we report for the first time that MICU2 is a guardian of mitochondrial respiratory chain function and a pivotal element during the cancer metabolic switch from oxidative metabolism (oxidative phosphorylation [OXPHOS]) to glycolysis. Moreover, our data suggest that the fine regulation of the MICU1–MICU1 and MICU2–MICU1 dimers in association with MCU may shape the major metabolic transformations in cancer.

## Results

### MICU2 expression is higher in stage IV CRC and metastases

A recent investigation demonstrated that the MCU complex can be linked to overall aggressiveness of CRC.^16^ In particular, using The Cancer Genome Atlas Colon Adenocarcinoma (TCGA-COAD) dataset, the authors observed that the expression of MICU1 and MICU2 are respectively down- and upregulated with stages. The stoichiometry of regulatory units of MCU complex, MICU1 and MICU2, have been shown to modulate the uniporter-mediated Ca^2+^ influx.^17^

Here, we focused on investigating the expression of MICU1, MICU2, and the MICU2/MICU1 ratio in colon tumors using the cohort published by TCGA. As described previously, we found that MICU2 expression is significantly upregulated in colon tumors of patients with stage IV CRC. MICU3 is not expressed in the CRC. Interestingly, the MICU2/MICU1 ratio is significantly upregulated in colon tumors of stage IV compared with tumors belonging to less aggressive stages (Figure S1A). In another cohort (GSE41252) encompassing primary tumors but also liver and lung metastatic CRC samples, we observed that the expression of MICU1 is downregulated while both MICU2 expression and the MICU2/MICU1 ratio are significantly upregulated in liver metastasis of CRC (Figure 1B and S1B). In addition, the expression of MCU complex players according to CRC stages was verified on TCGA data. Significantly, MCU, MICU1 and MCUb were negatively expressed while MICU2 was positively expressed (Figure 1B). Thus, the MICU2/MICU1 ratio is positively correlated to the aggressiveness of a CRC tumor, suggesting that the stoichiometry of the regulatory units of MCU complex may be critical for the tumor development.

**Figure 1.**
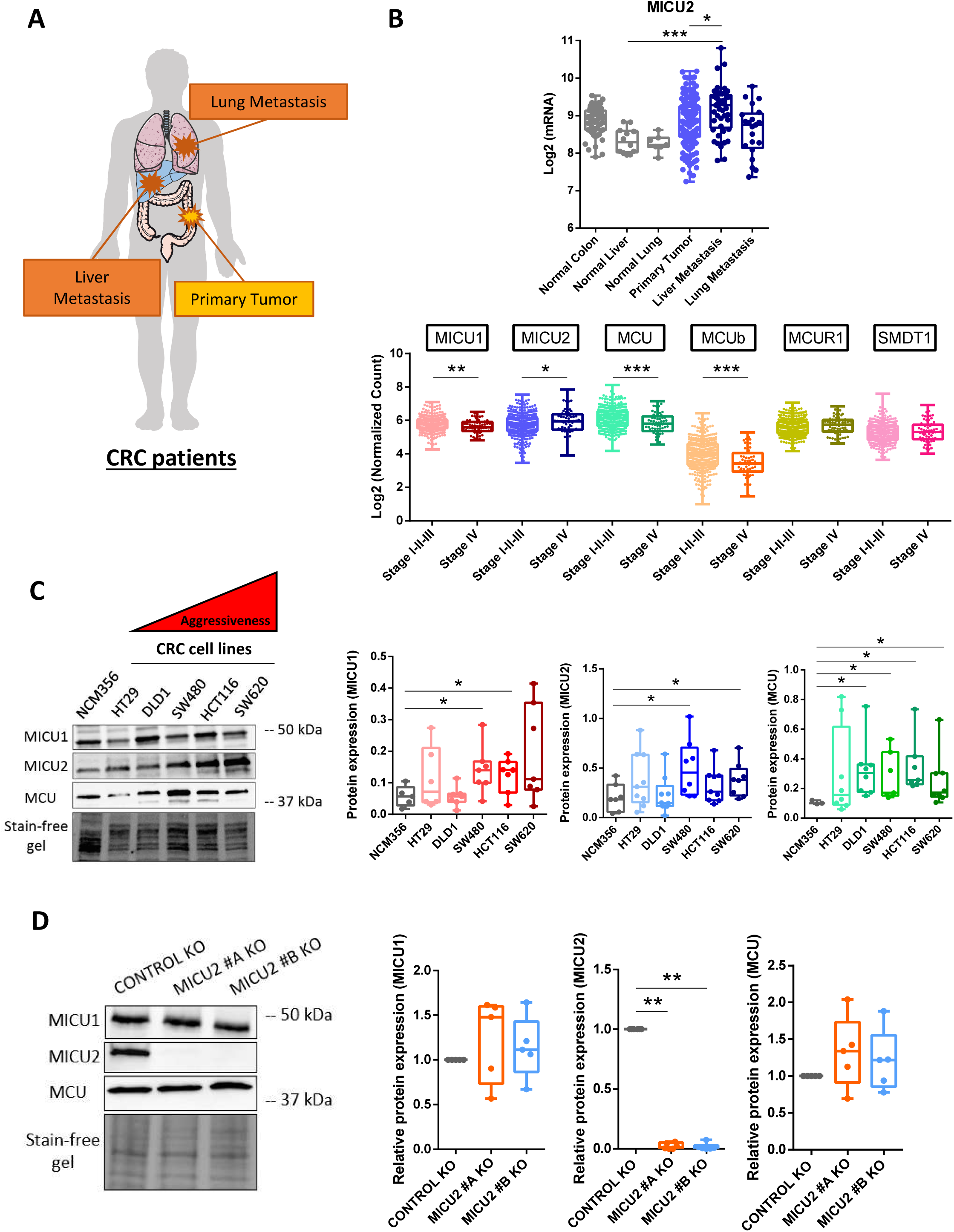
Expression of MICU2 in stage IV CRC and in metastases from patients with CRC. (A) Representative scheme of organs used for the bioinformatics analysis of normal, primary tumor, and metastatic samples. (B) Transcriptomic analysis of MICU2 in the GSE41258 (top panel) datasets and of the expression of the players in the MCU complex in the TCGA (bottom panel). Each data point represents an individual sample (ANOVA followed by Dunn’s multiple comparisons test). (C) Left panel: representative western blot of MCU, MICU1, and MICU2 in a panel of CRC cell lines with variable aggressiveness. Right panel: quantification of the protein expression of MCU, MICU1, MICU2 western blot (n=5–9, Mann–Whitney test). (D) Representative western blot of MCU, MICU1, and MICU2 expression in the HCT116 Control and MICU2 KO cell lines obtained by CRISPER-Cas9 methodology. Quantification of the protein expression of MICU1, MICU2, and MCU in the HCT116 Control KO, MICU2 #A KO, and MICU2 #B KO cell lines (n=6, Kruskal–Wallis test). On all plots, *p<0.05, **p<0.01, and ***p<0.001.

Using western blotting, we compared the protein expression of MCU, MICU1, and MICU2 in a non-tumor colorectal epithelial cell line, NCM356, and in five CRC cell lines (HT29, DLD1, SW480, HCT116, and SW620) with different mutation and aggressiveness profiles. The tumor cell lines showed significantly higher expression of MCU and MICU1 than the non-tumor cell line. Interestingly, MICU2 expression was significantly higher in the more aggressive tumor cell lines (SW480, HCT116, and SW620) compared with the less aggressive cell lines (HT29 and DLD1) and the non-tumor cell line (NCM356) (Figure 1C). As MICU2 cannot bind to the MCU complex without MICU1, we modified the HCT116 cell line with CRISPR-Cas9 technology to generate two MICU2 knockout (KO) clones (Figures 1D and S1C) to modulate the MICU2/MICU1 ratio. Deletion of MICU2 did not alter the protein expression of MCU and MICU1 (Figure 1D), but MICU1 messenger RNA (mRNA) expression was significantly reduced by 25% in one of the MICU2 KO clones (Figure S1C).

### MICU2 controls CRC cell proliferation *in vitro* and CRC tumor growth *in vivo*

Primary tumors of TCGA-COAD were classified as “High” or “Low” for MICU1, MICU2 and the ratio of MICU2/MICU1 if the value of their expression/ratio are respectively superior to the 95^th^ percentile or inferior to the 5^th^ percentile of the expression/ratio in normal samples (Figure S1D). Using this classification of tumors samples, we then performed a differential expression analysis (DEA) followed by a gene set enrichment analysis (GSEA) to identify genes and pathways that are differentially modulated based on the status of MICU2 or the MICU2/MICU1 ratio (Figure 2A and S2A). Interestingly, we observed that cell cycle–related gene sets are frequently enriched in tumor samples with high MICU2 expression and tend to be less represented in tumor samples with a low MICU2 expression (Figures 2B and S2B).

**Figure 2.**
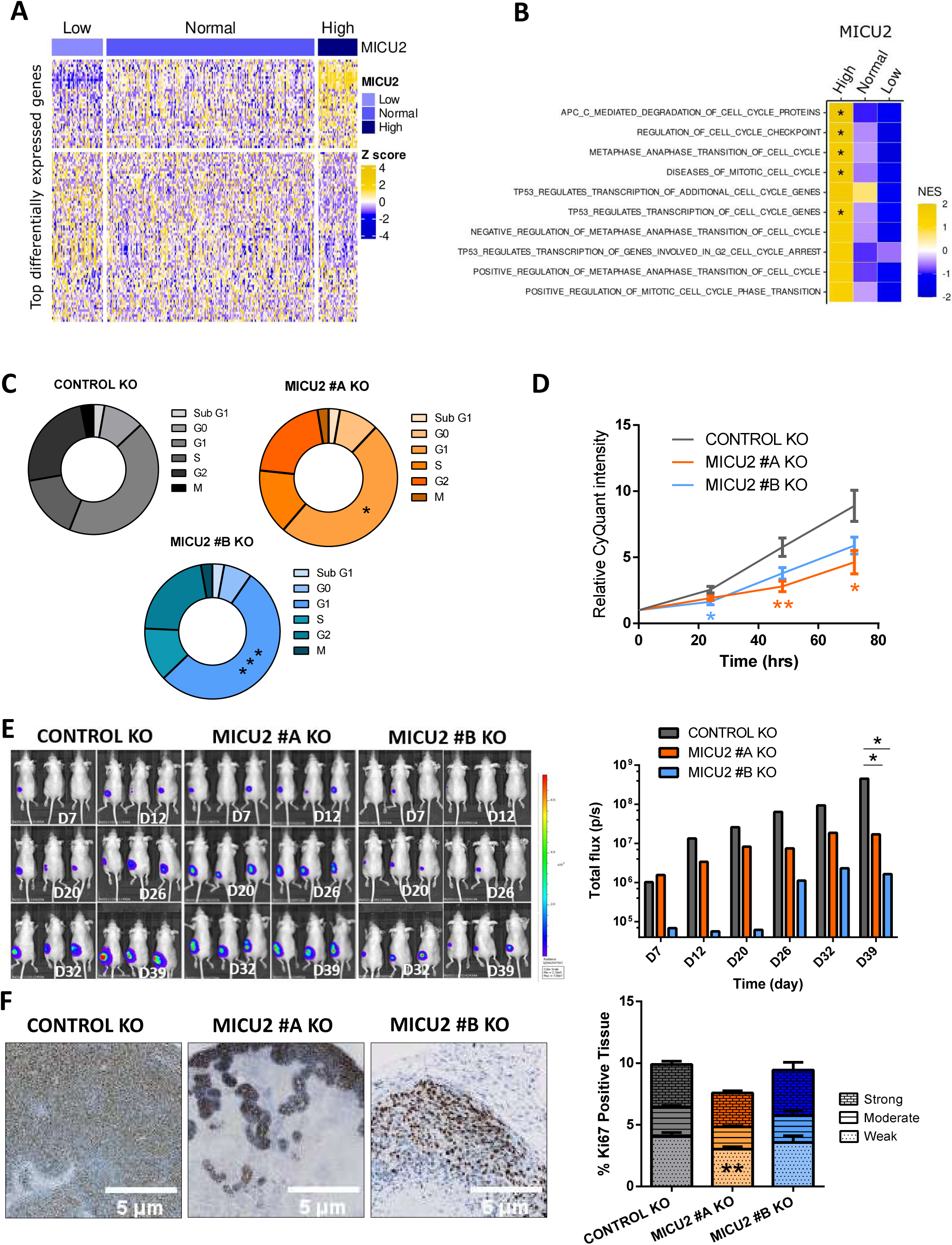
Involvement of MICU2 in the CRC tumor phenotype. (A) Heatmap of the most differentially expressed genes between colon tumor samples in the TCGA-COAD dataset with low, normal, or high expression of MICU2. B) Heatmap of normalized enrichment scores of the top 10 cell cycle–associated gene signatures obtained by GSEA for primary colon tumor samples with low, normal, or high expression of MICU2. (C) Graphical representation of the cell cycle in the Control and MICU2 KO cells lines (n=6, ANOVA followed by Dunnett’s multiple comparisons test). (D) CyQuant proliferation analysis of the Control and MICU2 KO cells lines (n=11, Kruskal– Wallis test). (E) Representative bioluminescence images of cancer progression and metastasis in mice injected with the luciferase-expressing Control or MICU2 KO cells lines and quantification of whole-body bioluminescence (n=10, ANOVA followed by Dunnett’s multiple comparisons test). (F) Representative images of tumoral tissue sections stained for Ki67 from xenografted Control and MICU2 KO cells lines and quantification of the percentage of tissue positive for Ki67. The intensity of positivity was scored as weak, moderate, or strong. The scale bar is 5µm (n=6, ANOVA followed by Sidak’s multiple comparisons test). On all plots, *p<0.05, **p<0.01, and ***p<0.001.

Cell cycle analysis by flow cytometry demonstrated an increase in the percentage of cells in the G1 phase associated with a decrease in proliferation of the MICU2 KO cell lines compared with the Control KO cell line (Figures 2C and 2D). The HCT116 control and MICU2 KO cell lines were engineered to endogenously express luciferase and injected subcutaneously into NOD-SCID mice. There was a significant reduction in total bioluminescence in mice injected with HCT116 MICU2 KO cells compared with mice injected with the control HCT116 cells (Figure 2E). To confirm the role of MICU2 in CRC tumor growth, we performed Ki67 staining on tumor sections. We noted a significant reduction in cell proliferation for MICU2 #A KO tumors but not for MICU2 #B KO tumors (Figure 2F). This outcome could be potentially explained by the delay we observed for the tumor engraftment of MICU2 #B KO cells. Hence, at the end of our experiment, these tumors were still in the exponential growth phase.

To further investigate the role of MICU2 in cancer, we investigated its role in the response to CRC chemotherapy treatment. We evaluated the dose-response of oxaliplatin and 5-fluorouracil (5FU) in the Control and MICU2 KO cell lines. Both cell lines showed a concentration-dependent reduction in cell viability, but without a significant difference in the sensitivity to these chemotherapeutic agents (Figure S2C). Based on the effect of MCU that has been described in cancer cells, we evaluated the role of MICU2 in cell motility (wound healing and spheroid migration assays). We noted decreased cell migration in the MICU2 KO cell lines compared with the Control KO cell line (Figures S2D and S2E).

Taken together, these results demonstrate that MICU2 expression is associated with several cancer cell properties. In particular, MICU2 is critical for cancer cell proliferation and the cell cycle. MICU2 KO can also reduce the migration of CRC cells.

### MICU2 protects CRC cells against fragmentation and controls mitoCa^2+^ uptake

Mitochondria share different contact sites with the endoplasmic reticulum (ER) and the plasma membrane.^18,19^ During the movement of Ca^2+^ from the extracellular or intracellular compartments, the mitoCa^2+^ uptake machinery shapes the cellular response. The mutual interplay between Ca^2+^ homeostasis and mitochondrial morphology appears to be an essential element in mitochondrial dynamics and Ca^2+^ signalling.^20^

To understand the impact of MICU2 KO on the mitochondrial morphology and network, we first evaluated the number of mitochondria and the amount of mitochondrial DNA in the Control and MICU2 KO cell lines. There were no differences in these two parameters between the cell lines (Figure 3A and 3B). We then assessed the network and size of the mitochondria using confocal imaging. Interestingly, MICU2 KO cell lines have a significant reduced number of mitochondria with an area >20 µM3 compared to the Control KO cell line. This suggest that there is a decrease in the mitochondrial connection network and therefore greater fragmentation (Figure 3C). To confirm this hypothesis we analysed proteins linked to mitochondrial dynamics, namely the expression of OPA1 and MFN2, which are involved in fusion, and the expression of DRP1 and its active phosphorylated form (P-DRP1 Ser616), which is involved in fission.21 -There was no difference in the expression of OPA1 or MFN2, but the P-DRP1/DRP ratio was increased in MICU2 KO cell lines, indicating increased mitochondrial fission, consistent with fragmentation of the mitochondrial network (Figure 3D). The mRNA expression of fission (DRP1 and FIS1) and fusion (OPA1 and MFN2) genes was not significantly altered in MICU2 KO cell lines compared to the Control KO cell line (Figure S3A).

**Figure 3.**
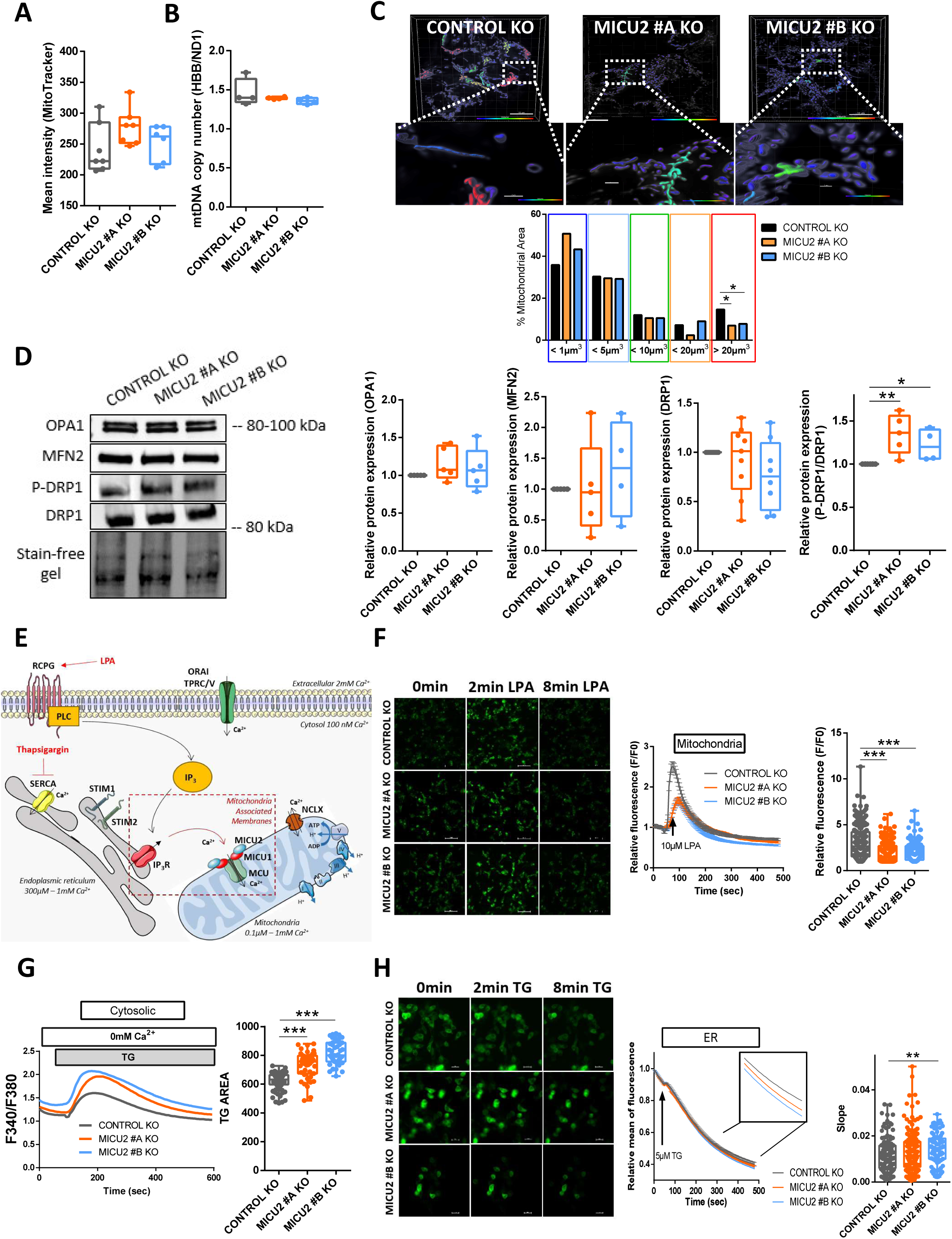
The role of MICU2 in the mitochondrial network and Ca^2+^ signaling. (A) Quantification of the number of mitochondria in the Control and MICU2 KO cell lines with Mitotracker green. (B) Quantification of mitochondrial DNA copy number by RT-qPCR using primers HBB and ND1. (C) Top panel: representative 3D images of the mitochondrial network of the HCT116 Control and MICU2 KO cell lines colored based on the mitochondrial volume. The enlargement shows an example of mitochondria with different shapes. The scale bar is 10 μm. Bottom panel: bar graphs representing the distribution of the mitochondrial population based on the mitochondrial volume. Anova followed by Dunnett’s multiple comparisons test. (D) Left panel: representative western blot of mitochondrial fusion (OPA1 and MFN2) and fission (DRP1 and P-DRP1) proteins in the Control and MICU2 KO cell lines. Right panel: relative protein expression of mitochondrial fusion and fission proteins (n=4–5, ANOVA followed by Dunn’s multiple comparisons test). (E) Schematic illustration of LPA-induced or TG-induced cytosolic Ca^2+^ and mitoCa^2+^ signaling in non-excitable cells. (F) Left panel: representative images of the Control and MICU2 KO cell lines transfected with the genetically encoded mitoCa^2+^ indicator mt-riG6m and stimulated with LPA. The scale bar is 100 µm. Right panel: relative Ca^2+^ responses induced by LPA in Control and MICU2 KO cell lines expressing the mitoCa^2+^ probe mt-riG6m (the data are presented as the mean ± SEM of 8 independent experiments regrouping 354–594 cells, Kruskal–Wallis test). (G) Left panel: representative SOCE traces in the Control and MICU2 KO cell lines. Right panel: boxplot representing the ER-Ca^2+^ store depletion by TG in the absence of extracellular Ca^2+^ (n=6, N=42–95, ANOVA followed by Dunnett’s multiple comparisons test). (H) Left panel: representative images of the Control and MICU2 KO cell lines transfected with the genetically encoded ER Ca^2+^ indicator miGer and stimulated with TG. The scale bar is 20 µm. Right panel: relative ER Ca^2+^ content. The boxplot represents the slope of TG-induced ER Ca^2+^ release (n=6– 8, N=88–156, Mann–Whitney test). On all plots, *p<0.05, **p<0.01, and ***p<0.001.

MICU2 modulates mitoCa^2+^ uptake. Thus, we examined the consequences of the genetic deletion of MICU2 on mitochondrial, cytosolic, and ER Ca^2+^ signaling. Stimulation of a G protein–coupled receptor (GPCR) such as the lysophosphatidic acid (LPA) receptor by LPA induces inositol trisphosphate (IP_3_) production. IP_3_ then activate IP_3_ receptors leading to a Ca^2+^ release from ER and induce mitochondria Ca^2+^ influx through the uniporter MCU (Figure 3E).

We assessed mitoCa^2+^ homeostasis by using a genetically encoded Ca^2+^ indicator targeting the mitochondrial compartment (mt-riG6m). Stimulation of cells with LPA induced a transient increase in mitoCa^2+^ that was significantly reduced in the MICU2 KO cell lines compared with the Control KO cell line (Figure 3F). We then wondered whether this reduction in mitoCa^2+^ influx might affect cytosolic Ca^2+^ homeostasis. Stimulation of transitory ER Ca^2+^ release by LPA activates store-operated calcium entry (SOCE).^21^ MICU2 KO had no effect on SOCE induced by LPA (Figure S3B). Interestingly, thapsigargin (TG)-mediated release of ER Ca^2+^ stores were increased in the MICU2 KO cell lines compared with the Control KO cell line without modifying SOCE (Figures 3G and S3C). We then assessed whether this change is due to higher Ca^2+^ content in ER stores. Using the genetically encoded Ca^2+^ indicator targeting ER (miGer), we observed a small but significant increase in the speed of the TG-induced ER Ca^2+^ release in the MICU2 KO #B cell line (Figure 3H). Using an intracellular Ca^2+^ chelator (BAPTA-AM), we assessed the viability of the Control KO and MICU2 KO cell lines. Treatment of the cells with BAPTA-AM resulted in a significant decrease in the viability of the Control KO and MICU2 KO cell lines (Figure S3D). Altogether, these results suggest that genetic deletion of MICU2 impairs mitoCa^2+^ homeostasis and the ability of mitochondria to buffer variations in cytosolic [Ca^2+^].

Crosstalk between Ca^2+^ and reactive oxygen species (ROS) is found in many pathologies including cancers.^22^ Hence, we measured ROS production in HCT116 Control and MICU2 KO clones. We analyzed the levels of total superoxide ions (DHE) and mitochondrial superoxide (MitoSox) and the amount of hydrogen peroxide (DCFDA). We observed no difference between the Control and MICU2 KO cell lines (Figure S3E). In addition, there was no difference in the viability of the Control and MICU2 KO cell lines treated with a mitochondria-targeted antioxidant (MITOTEMPO) (Figure S3F). This outcome confirms that genetic deletion of MICU2 does not affect ROS production.

Taken together, these results reveal that MICU2 is necessary for the stability and the quality of mitochondrial network while allowing the proper management of ER-mitochondria Ca^2+^ flux.

### Deletion of MICU2 reduces respiratory complex expression and activity associated with loss of respiratory chain sensitivity to Ca^2+^

MitoCa^2+^ plays critical signaling roles in regulating ATP production linked to oxygen consumption by mitochondria. We hypothesized that the deletion of MICU2 alters the functioning of the respiratory chain.

First, we tested this hypothesis by analyzing in TCGA-COAD dataset based on the expression of MICU1, MICU2, and the MICU2/MICU1 ratio. The mRNA expression of genes belonging to the mitochondrial respiratory chain complexes are not modified by the expression of MICU1, MICU2, or the MICU2/MICU1 ratio (Figure S4A). However, we observed that TCGA-COAD tumor samples with a “high” MICU1 status show higher expression of genes associated with the assembly of complexes I and IV of the mitochondrial respiratory chain compared with tumor samples with a “low” MICU1 status (Figure 4B and S4B). In contrast, we observed that TCGA-COAD tumor samples with a “high” status of MICU2 and the MICU2/MICU1 ratio present higher expression of genes associated with the assembly of complexes of the mitochondrial respiratory chain compared with tumor samples with a “low” status for MICU2 and the MICU2/MICU1 ratio (Figure 4B and S4B), suggesting that the expression of MICU2 may contribute to the assembly complexes or supercomplexes. To validate this finding, we analyzed their expression and assembly into supercomplexes by Blue-Native PAGE. We observed a reduction in assembly of the I_1_III_2_IV supercomplex related to a significant reduction in the quantity of complex IV (Figure 4C). Consistently, the maximal activity of complex IV (cytochrome *c* oxidase) in the MICU2 KO cell lines was decreased compared with the Control KO cell line (Figure 4D). Mitochondria is the main organelle supporting energy supply of cells.^23^ The reduced expression of the terminal enzyme of the respiratory chain, complex IV, in MICU2 KO cells could limit the mitochondrial oxidative capacity. Inhibition of complex I and ATP synthase with rotenone and oligomycin, respectively, had a more pronounced effect on the viability of the Control KO cell line than the MICU2 KO cell lines (Figure 4E), suggesting that the MICU2 KO cell line is less dependent on oxidative phosphorylation (OXPHOS). In parallel, we verified that oligomycin and rotenone treatment did not significantly modify the mRNA expression of MICU1, MICU2, or MCU (Figure S4C).

**Figure 4.**
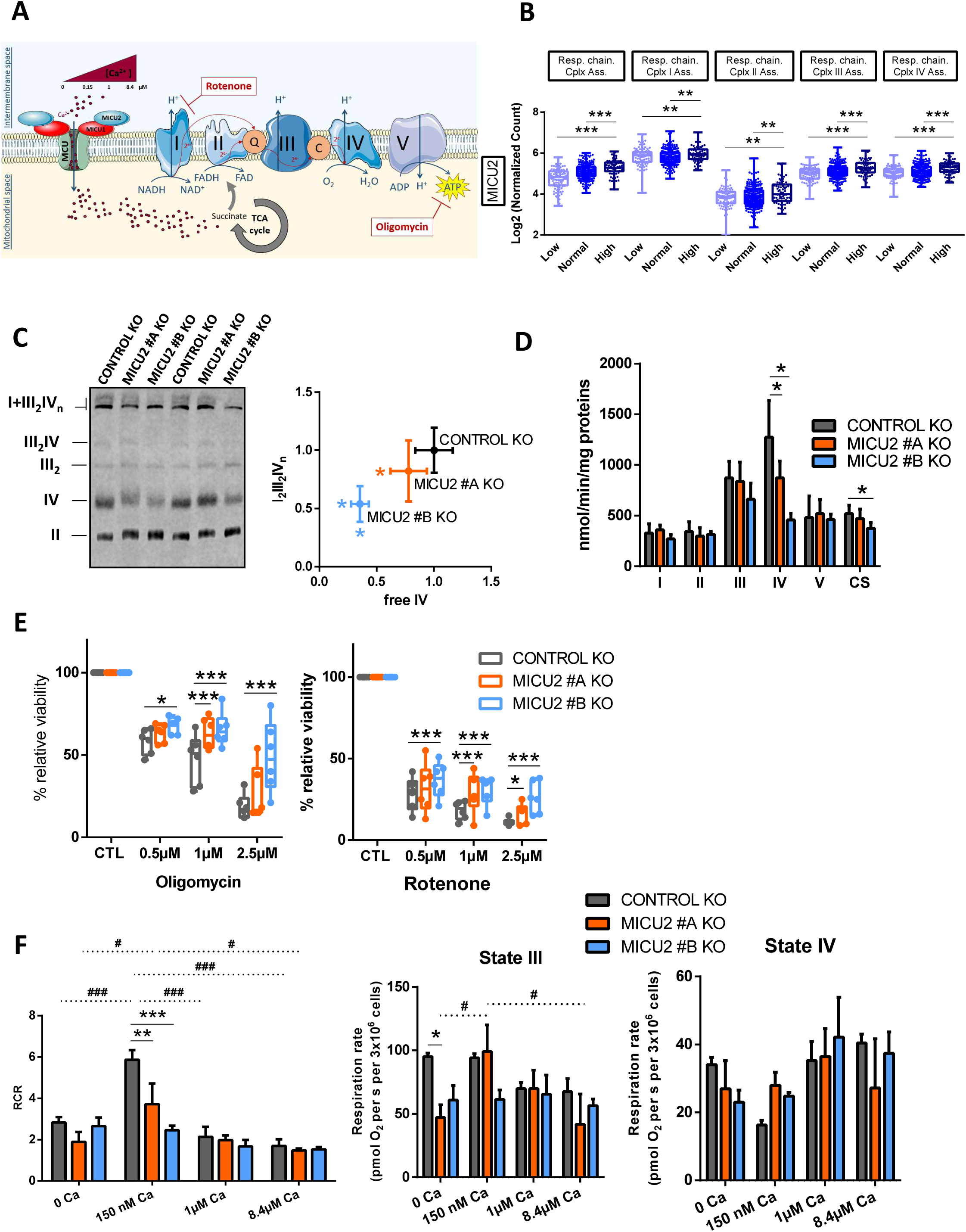
Contribution of MICU2 to respiratory chain complexes. (A) Simplified diagram of the respiratory chain and ATP synthesis in the mitochondria by OXPHOS. The production of ATP is possible due to the formation of a proton gradient around the membrane generated by the energy of the electrons supplied by NADH and FADH_2_ to this chain. These electrons are transported via different complexes: I, II, III, and IV. This diagram also illustrates the analysis of mitochondrial cellular respiration of different [Ca^2+^] (0, 150nM, 1µM, and 8.4µM). Ca^2+^ can regulate succinate, which in turn regulates complex II. (B) Boxplot representing the average expression of genes associated with the assembly of mitochondrial respiratory chain complexes I, II, III, and IV based on the status MICU2 in primary colon tumor samples from the TCGA-COAD dataset. (C) Analysis of supercomplex and complex assembly through BN-PAGE followed by western blotting. Left panel: representative blot hybridized with anti-NDUFS2 (complex I), UQCRC2 (complex III), SDHA (complex II), and MT-CO1 (complex IV). The position of the different free complexes or supercomplexes are indicated. Right panel: the supercomplex I1III2IVn quantity normalized to the CII quantity (y-axis) according to the free CIV quantity, normalized to the CII quantity (x-axis). The data are presented as the mean ± standard deviation (n=6, Mann–Whitney test). (D) Maximal activities of the respiratory chain complexes I (NADH ubiquinone reductase), II (succinate ubiquinone reductase), III (ubiquinol cytochrome *c* reductase), and complex IV (cytochrome oxidase), and activity of complex V (F1-ATPase) and the TCA enzyme CS. The data are presented as the mean ± standard deviation (n=5, Mann–Whitney test). (E) Boxplots representing the viability of the Control and MICU2 KO cell lines cultured in the presence of oligomycin (complex V inhibitor) (left panel) or rotenone (complex I inhibitor) (right panel) at increasing doses (0.5, 1, and 2.5µM) for 48 h (n=6, ANOVA followed by Dunnett’s multiple comparisons test). (F) Barplots representing respiratory states III, IV and RCR defined by the difference in oxygen consumption between succinate and substrate-free respiration (state II), phosphorylating respiration measured after injection of ADP and succinate (state III), and oxygen consumption after injection of oligomycin (state IV). The RCR is the respiratory control ratio calculated by the ratio between state III and IV. Respiratory rates are plotted against different [Ca2+] levels (0, 150nM, 1µM, and 8.4µM) (n=6-9, ANOVA followed by Dunnett’s multiple comparisons test). On all graphs, *p<0.05, **p<0.01, and ***p<0.001.

Variation in the mitoCa^2+^ concentration can have a dual role on OXPHOS activity. First, it can directly modify the activity of respiratory complexes. Second, Ca^2+^ can activate the tricarboxylic acid (TCA) cycle by affecting isocitrate dehydrogenase and α-ketoglutarate dehydrogenase, thus increasing OXPHOS. Using permeabilized cells, we evaluated the consequences of different cytosolic [Ca^2+^]—0, 150nM, 1µM, and 8.4µM—on OXPHOS stimulated by succinate, to avoid the consequence of Ca^2+^ variation on the activity of TCA cycle and focus only on the role of Ca^2+^ on the respiratory chain.^24^ We were particularly interested in state III of mitochondrial respiration stimulated by adenosine diphosphate (ADP) and state IV which is the inhibition of ATP synthase mainly controlled by proton leakage in the presence of substrate, so we defined the respiratory chain ratio (RCR) as an indicator of proper functioning of the respiratory chain.^25^

Globally, no significant effect of either the variation of the cytosolic [Ca^2+^] or the deletion of MICU2 is observed on both state III or state IV. However, the optimal function of the respiratory control ratio (RCR) is observed at physiological cytosolic [Ca^2+^] (150nM). At this concentration, the deletion of MICU2 significantly impairs the RCR. In absence of cytosolic [Ca^2+^] or at supra-physiological cytosolic [Ca^2+^], we observe a reduced RCR and no significant effect of the deletion of MICU2 on the RCR. MICU2 optimises mitochondrial respiration under physiological [Ca^2+^].

Overall, we demonstrated that MICU2, unlike MICU1, controls the complex biogenesis and activity of the respiratory chain and its presence optimizes oxygen consumption linked to mitochondrial ATP production in an environment with a physiological cytosolic [Ca^2+^]. We therefore took a closer look at metabolic rewiring and the substrate supply pathways regulated by calcium.

### MICU2 controls proliferation of CRC cells by promoting fatty acid oxidation and mitochondrial pyruvate utilization

Therefore, we studied the cellular respiration of intact Control and MICU2 KO cells, in standard RPMI medium with the presence of glucose (2g/L) and glutamine (0.4g/L). We observed a significant reduction in the basal respiration rate (oxidative metabolism), the respiration dedicated to ATP production–linked oxygen consumption, and the maximal respiration (maximal oxidative capacity) in the MICU2 KO cell lines compared with the Control KO cell line (Figure 5A), which could be linked to complex IV deficiency. However, mitoCa^2+^ allows adjusting ATP production–linked oxygen consumption to the cellular energy demand by directly or indirectly regulating PDH, isocitrate dehydrogenase, and α-ketoglutarate dehydrogenase activities. Thus, the decreased oxidative metabolism and mitochondrial ATP synthesis could also be due to limited substrate supply. We addressed the contribution of MICU2 to the preferred pathways and substrates (glucose, glutamine, and fatty acids) used for energy production by the mitochondria. Then we evaluated the importance of the three metabolic pathways separately.

**Figure 5.**
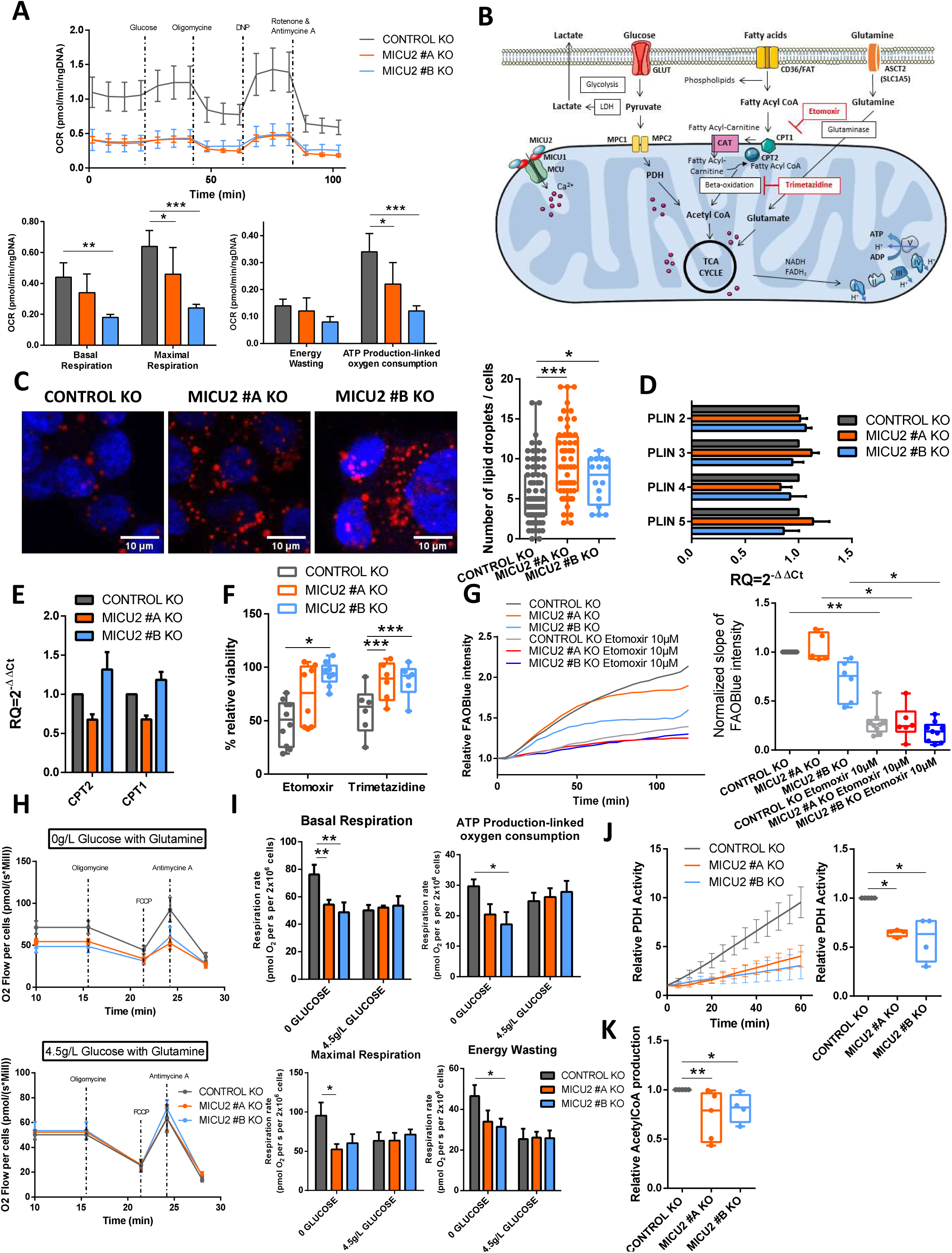
MICU2 modulates substrate selection for the TCA cycle. (A) Top panel: graph representing the OCR as a function of time in the Control and MICU2 KO cell lines and with sequential addition of glucose, oligomycin, DNP (protonophore), antimycin A (complex III inhibitor), and rotenone. Bottom panel: bar plots representing the basal respiration, maximal respiration, proton leak, and ATP production in Control and MICU2 KO cell lines (n=5, ANOVA followed Sidak’s multiple comparisons test). (B) Illustration of the different metabolic pathways essential for proper functioning of the mitochondria through the TCA cycle. (C) Left panel: representative images of lipid droplets (red) and nuclei (blue) in the Control and MICU2 KO cell lines. The scale bar is 10 µm. Right panel: boxplot representing the number of lipid droplets per cell (n=17– 80 cells, Mann–Whitney test). (D) qRT-PCR data showing fold changes in mRNA levels of fatty acid degradation proteins in the Control and MICU2 KO cell lines (n=8). (E) qRT-PCR data showing fold changes in mRNA levels of carnitine palmitoyltransferase 1 and 2 (CPT1 and CPT2) proteins in the Control and MICU2 KO cell lines. The results are expressed as the mean ± SEM of four and five independent experiments. (F) Boxplot representing the viability of the Control and MICU2 KO cell lines cultured for 48 h in the presence of 10 µM etomoxir (CPT1a inhibitor) or 10 µM trimetazidine (beta-oxidation inhibitor) (n=6–9, ANOVA followed by Dunnett’s multiple comparisons test). (G) Left panel: representative curve of FAOBlue fluorescence relative intensity in the Control and MICU2 KO cell lines with or without 10µM etomoxir. Right panel: boxplot-representing slope of FAOBlue fluorescence relative intensity with or without etomoxir (10µM). The data are presented as the mean ± standard error (n=5-9, Kruskal-Wallis test). (H) Graph representing the oxygen consumption rates of the Control and MICU2 KO cell lines in the absence with glutamine (top panel) or in the presence of high glucose (4.5g/L) and glutamine (bottom panel) in basal conditions and with sequential addition of oligomycin, FCCP (uncoupler) and antimycin A (complex III inhibitor). (I) Bar plots representing the basal respiration and ATP production–linked oxygen consumption (top panel) with maximal respiration and energy wasting (bottom panel) in the Control and MICU2 KO cell lines in presence of glutamine with or without high glucose (4.5g/L). The data are presented as the mean ± standard error of the mean (n=6, ANOVA followed Dunnett’s multiple comparisons test). (J) Left panel: PDH activity at 60min in the Control and MICU2 KO cell lines. Right panel: boxplot representing the PDH in the Control and MICU2 KO cell lines (n=4, ANOVA followed by Dunn’s multiple comparisons test). (K) Boxplot representing the production of acetyl-CoA in RPMI culture medium in the Control and MICU2 KO cell lines (n=4– 5, Kruskal–Wallis test followed by Dunn’s multiple comparisons test). On all plots, *p<0.05, **p<0.01, and ***p<0.001.

Fatty acids are transported into the cytosol and mitochondria by CD36 and carnitine palmitoyltransferase 1 (CPT1)/CPT2, respectively. Once inside the mitochondria, fatty acids are degraded by beta-oxidation to produce, among other things, acetyl-CoA that fuels the TCA cycle and energy production (Figure 5B). Here, using Oil Red O staining, we observed an increase in lipid droplets in the MICU2 KO cell lines compared with the Control KO cell line, suggesting either greater fatty acid storage or reduced beta-oxidation (Figure 5C). There was no difference between the Control and MICU2 KO cell lines in the mRNA expression of perilipins (PLIN), proteins that assemble on the surface of lipid droplets, or of CPT1 and CPT2 (Figures 5D and 5E). Compared with the Control KO cell line, the viability of the MICU2 KO cell lines was not reduced as much by the inhibitors of fatty acid CoA transporters CPT1 and CPT2 (etomoxir) or beta-oxidation (trimetazidine). There was no significant difference in the mRNA expression of MCU, MICU1, and MICU2 after etomoxir or trimetazidine treatment (Figures 5F and S5A). This suggests that in the absence of MICU2, the HCT116 cell line tends to be less dependent of fatty acid metabolism.

We then wondered whether the mitochondrial energy production mediated by fatty acid utilization depends on MICU2 expression. Hence, we used a fluorescent indicator of fatty acid oxidation (FAOBlue), which indicated that the MICU2 KO #B cell line had reduced fatty acid oxidation compared with the Control and MICU2 KO #A cell lines (Figure 5G).

Taken together, these results suggest that fatty acid beta-oxidation is not the preferred energy pathway used by MICU2 KO cells and that the deletion of MICU2 could alter the quantity of lipid droplets and beta-oxidation.

Glutamine is transformed into glutamate to fuel the TCA cycle. Glutaminolysis is widely used in HCT116 cells to sustain TCA cycle anaplerosis.^26^ To determine whether MICU2 expression regulate the oxidative metabolism sustain through glutamine oxidation, we first analyzed respiration rates on intact cells in the only presence of glutamine (Figure 5H). With glutamine as substrate, basal respiration rate, respiration-linked to ATP synthesis and maximal oxidative capacity were all significantly reduced in MICU2 KO cells. However, we did not note significant difference in the decrease of the viability of Control and MICU2 KO cells induced by inhibitors of the amino acid transporter (V-9302) and glutaminase (telaglenastat) (Figure S5B). Glutamine has a versatile role in cell metabolism, not only participating in TCA cycle anaplerosis but also the biosynthesis of nucleotides, glutathione (GSH), and other nonessential amino acids. Thus, this suggests that MICU2 KO cells still displayed dependency on glutamine metabolism for their survival, despite a reduced mitochondrial oxidative metabolism support by glutamine.

High glucose concentration (4.5g/l) is known to force cellular metabolism towards “anaerobic” glycolysis. Indeed, the addition of high glucose decreases oxidative metabolism (basal respiration, ∼80 nmolO2/sec/million cells in glutamine versus ∼55nmolO2/sec/million cells in glutamine and high glucose condition, Figures 5H and 5I) in control cells but has no effect on MICU 2KO cells, suggesting that these cells already have high reliance on glycolytic metabolism.

We thus measured PDH activity (known to be regulated by Ca^2+^) and acetyl-CoA production and observed a significant reduction in the MICU2 KO cell lines compared with the Control KO cell line (Figures 5J and 5K). We observed a significant increase in PDP1 and PDHA1 mRNA expression in the MICU2 KO cell lines (Figures S5C and S5D), suggesting an increase in the expression of pyruvate regulator alongside a decrease in global PDH activity. To take our analysis a step further, we used mRNA expression to study the pentose phosphate pathway. PRPS1 and PRPS2 were increased in MICU2 KO cell lines (Figure S5E). Jing et al.^27^ showed that PRPS1 upregulation is more important in promoting tumorigenesis and is a promising diagnostic indicator of CRC.

Overall, our results indicate that the MICU2 KO cell lines have lower OXPHOS associated with a reduction in PDH activity and acetyl-CoA production. Interestingly, the balance between glucose, beta-oxidation and glutaminolysis is modified in MICU2 KO cells.

### MICU2 knockout cells exhibit increased glycolytic flux *in vitro* and *in vivo*

Cellular ATP production is dependent of the TCA cycle and glycolysis, both of which are associated with glucose uptake by GLUT transporters. In the cytosol, glycolysis transforms glucose into pyruvate while producing ATP. Then, pyruvate is degraded into lactate by lactate dehydrogenase (LDH) and excreted from the cell by lactate transporters (MCT). Pyruvate can be transported into mitochondria through MPC proteins. PDH produces acetyl-CoA that serves as substrate for TCA cycle and mitochondrial ATP production (Figure 6A).

**Figure 6.**
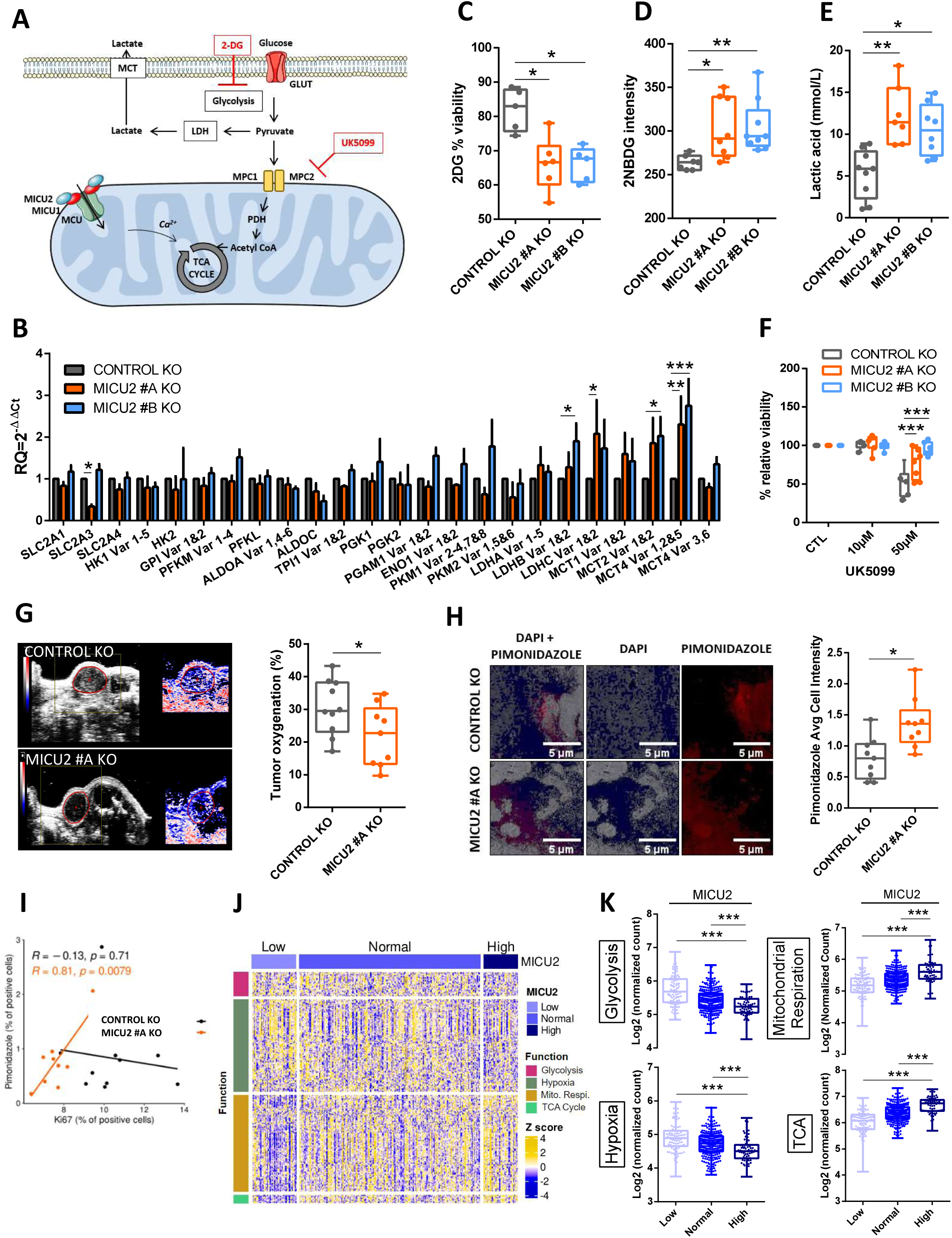
MICU2 expression correlates negatively with glycolysis and hypoxia and is associated with OXPHOS and the TCA cycle. (A) Diagram showing the glycolytic pathway. Glucose crosses the plasma membrane via GLUT; pyruvate is then reduced to lactate or can also pass the mitochondrial membrane to produce acetyl-CoA via PDH. Inhibitors of glycolysis such as 2DG (a non-metabolizable glucose analog) and UK5099 are present at their target. (B) Bar plots representing the level of expression of glycolysis-related genes in the Control and MICU2 KO cell lines (n=3–13, ANOVA followed Dunnett’s multiple comparisons test). The data are presented as the mean ± SEM. (C) Boxplot representing viability of the Control and MICU2 KO cell lines after culturing for 48h with 2DG (n=5–6, ANOVA followed by Dunn’s multiple comparisons test). (D) Boxplot representing the fluorescence intensity of the Control and MICU2 KO cell lines incubated for 30 min with the fluorescent glucose analog 2NBDG (n=7–9, ANOVA followed by Dunn’s multiple comparisons test). (E) Boxplot representing lactic acid production in the Control and MICU2 KO cell lines (n=7–9, ANOVA followed by Dunn’s multiple comparisons test). (F) Boxplot representing viability of the Control and MICU2 KO cell lines cultured in the presence of UK5099 (MPC inhibitor) (n=6–9, Anova followed by Dunnett’s multiple comparisons test). (G) Left panel: representative images of photoacoustic imaging in the Control and MICU2 KO cell lines illustrating the oxygen levels in tumors at day 32. Right panel: boxplot representing tumor oxygenation of the Control and MICU2 KO cell lines (n=9–10, Mann–Whitney test). (H) Left panel: representative images of hypoxic areas in tumors detected by pimonidazole (red) and DAPI (blue) staining. The scale bar is 5µm. Right panel: boxplot representing the quantification of pimonidazole-positive areas on whole tumor sections (n=9, Mann–Whitney test). (I) Scatter plot representing the correlation of the Ki67 labeling index versus the pimonidazole score in the Control and MICU2 KO cell lines (R is the Pearson correlation coefficient). (J) Heatmap representing the expression of genes associated with glycolysis, hypoxia, mitochondrial respiration, and the TCA cycle as a function of the MICU2 status of primary tumor samples from the TCGA-COAD dataset. (K) Boxplot representing the mean expression of genes associated with glycolysis, hypoxia, mitochondrial respiration, and the TCA cycle according to the MICU2 status of primary tumor samples of the TCGA-COAD dataset. On all plots, *p<0.05, **p<0.01, and ***p<0.001.

Here, we observed significantly increased mRNA expression of LDH and lactate transporters (MCT2 and MCT4) in the MICU2 KO cell lines compared with the Control cell line (Figure 6B), suggesting a more pronounced dependence on anaerobic glycolysis in these cell lines. Consistently, the MICU2 KO cell lines showed reduced viability when cultured in presence of a non-metabolizable glucose analog (2-deoxy-D-glucose [2DG]) compared with only a modest effect for the Control KO cell line (Figure 6C). Incorporation of fluorescent glucose (2-NBDG) and the production of lactic acid were also higher in the MICU2 KO cell lines compared with the Control KO cell line (Figure 6D and 6E).

To determine the preference of MICU2 KO cells for a glycolytic phenotype, we treated them with UK5099 (MPC1/MPC2 pyruvate transporter inhibitor). In these conditions, pyruvate cannot be used to sustain oxidative metabolism but is mainly converted to lactate. The Control KO cell line was more sensitive to UK5099 than the MICU2 KO cell lines (Figure 6F). In parallel, we found that UK5099 treatment did not significantly modify the mRNA expression of MICU1, MICU2, or MCU (Figure S6A). This finding suggests that the MICU2 KO cell lines are less dependent on energy generated from pyruvate by mitochondria.

Cancer cells growing in a hypoxic environment tend to be more dependent on anaerobic glycolysis for energy production.^28^ As the MICU2 KO cell lines develop a preference for glycolysis, we measured the oxygenation of subcutaneous transplanted MICU2 KO tumors using photoacoustic imaging. We observed a significant reduction in the oxygenation of MICU2 KO #A tumors compared with Control KO tumors, suggesting a more hypoxic environment in those tumors (Figure 6G). This was confirmed by the staining of tumor sections with pimonidazole, a hypoxia indicator (Figure 6H). Interestingly, we found a positive correlation between the proliferative index (Ki67) and the pimonidazole only for MICU2 KO #A tumors, suggesting that the growth of MICU2 KO tumors is dependent on hypoxia (Figure 6I).

From the TCGA-COAD dataset, we defined three gene sets with genes associated with glycolysis, hypoxia, mitochondrial respiration, or the TCA cycle (Figures 6J and S6B). Interestingly, the expression of MICU1 is positively correlated with the average expression of genes associated with glycolysis (Figures S6C). By contrast, MICU2 expression and the MICU2/MICU1 ratio are negatively correlated with gene sets associated with glycolysis and hypoxia and positively correlated with gene sets associated with mitochondrial respiration and the TCA cycle (Figures 6K). Using the Cancer Cell Line Encyclopedia (CCLE) dataset, we also defined the expression of MICU1, MICU2, and the MICU2/MICU1 ratio in 57 colorectal cancer cell lines using the four gene sets. We observed a tendency for a positive correlation between the expression of MICU1 and the average expression of genes associated with glycolysis. By contrast, we noted a tendency for a positive correlation between the expression of MICU2 and the mean expression of genes associated with the TCA cycle. We found that the MICU2/MICU1 ratio tends to be negatively correlated with the mean expression of genes associated with glycolysis (Figures S6D and S6E). Thus, MICU2 expression seems to promote the expression of genes associated with mitochondrial energy production. Our results suggest that CRC cells with low MICU2 expression develop a preference for glycolysis to support cancer cell proliferation in a hypoxic environment.

## Discussion

In this study, we demonstrated that MICU2 is a guardian of mitochondrial OXPHOS and a pivotal element during the cancer metabolic switch from oxidative metabolism to glycolysis. The MCU complex mediates mitoCa^2+^. Within this complex a scaffold (protein) EMRE and mtCa^2+^ proteins (MICU1, MICU2, and MICU3) interact with MCU, a pore-forming unit able to modulate mitoCa^2+^ absorption. Deregulation of mitoCa^2+^ homeostasis has been described in several cancers. Consistently, MCU expression is elevated in various types of cancer, including breast cancer, hepatocellular carcinoma, melanoma and CRC.^16^

The Ca^2+^-dependent activation of MCU is conferred by MICUs, a set of EF hand–containing regulatory proteins. In all tissues this process is mediated by MICU1 and MICU2: MICU1 is responsible for keeping MCU in a closed state^29^ and MICU2 allows Ca^2+^-dependent MCU opening.^15^ In brain and muscle, MICU3 replaces MICU2 for this task.^30^ Several studies have endeavored to decipher molecular mechanism by which MICU1–MICU2 and MICU1–MICU3 heterodimers or MICU1–MICU1 homodimers regulate MCU channeling. However, the study of MICU proteins in a cancerous context remains very limited. In ovarian cancer, MICU1 silencing inhibits clonal growth, migration, and invasion *in vitro*, whereas its silencing *in vivo* inhibits tumor growth and increases cisplatin efficacy and overall survival.^31^ Interestingly, inhibition of MICU1 results in the concomitant disappearance of MICU2 protein expression, indicating that MICU2 requires MICU1 for its stability, but not vice versa.^15^ Thus, we investigated the stoichiometry of MICU2/MICU1 and the molecular function of MICU2 in cancer cells by knocking down the MICU2 gene in CRC cells. Bioinformatics analysis of publicly available CRC datasets revealed that transcriptomic expression of MICU2 or the MICU2/MICU1 ratio is correlated with the aggressiveness of CRC, with the highest expression measured in stage IV tumors and in metastases of CRC. These findings confirmed a previous report from Zhu et al.,^32^ who described poorer overall survival in patients with CRC and higher MICU2 expression. The authors proposed that expression of the different members of MCU complex could be associated with a risk signature that predicts the prognosis and responses to immunotherapy in patients with CRC.

We have provided the first (to the best of our knowledge) experimental evidence that MICU2-mediated mitoCa^2+^ uptake is crucial for CRC cell growth by stabilizing mitochondrial network (fusion/fission balance), respiratory chain complexes, OXPHOS and regulating main substrate supply pathways in CRC cells. These mechanistic data validate the bioinformatics analysis, which shows that the stoichiometry of the MICU2/MICU1 ratio and in particular the expression of MICU2 are correlated with the expression of genes related to mitochondrial respiration and the TCA cycle and inversely correlated with the expression of genes related to glycolysis in tumor samples.

### MitoCa^2+^ and mitochondria dynamics

We demonstrated that a loss of MICU2 decreases mitoCa^2+^ influx in CRC cells. Recently, Vishnu et al.^33^ described an equivalent role of MICU2 in mitoCa^2+^ uptake in β cells. The absence of MICU2 has been associated with reduced mitoCa^2+^ uptake, but it also leads to a decrease in the quantity of MCU.^13^ In CRC, mitoCa^2+^ uptake promotes TFAM dephosphorylation to enhance mitochondrial biogenesis.^34^ Changes in mitochondrial size and shape have been implicated in the regulation of mtCa^2+^ uptake. Using MNF2 KO or a DRP1 dominant negative mutation, Kowaltowski et al.^35^ described significant effects of these proteins: They are partially responsible for the effect of mitochondrial morphology on mitoCa^2+^ uptake but do not affect MCU expression. MFN2 KO decreases both mitoCa^2+^ uptake and Ca^2+^ retention capacity in mitochondria. DRP1 dominant negative mutation enhances both mitoCa^2+^ uptake capacity and rates. In mice, constitutive DRP1 ablation reduces muscle growth.^36^ In the present study, MICU2 KO did not affect the number of mitochondria, but the phospho-DRP1/DRP1 ratio was strongly increased and associated with a higher fragmentation level of mitochondria and reduced mitoCa^2+^ uptake. We demonstrated that MICU2 is an important actor for the regulation of mitoCa^2+^ uptake and mitochondrial morphology by altering phospho-DRP1/DRP1 activity.

### MICU2 promotes the functioning of the respiratory chain

MICU1 and MICU2 KO cells exhibit altered [Ca^2+^] threshold for mitoCa^2+^ uptake.^37^ MICU2 acts as the genuine gatekeeper of MCU at low cytosolic [Ca^2+^],^15^ despite the huge driving force for matrix cation entry, thus preventing mitoCa^2+^ accumulation and Ca^2+^ overload.^38^ MitoCa^2+^ acts as a key regulator of oxidative metabolism in mammalian cells.^39^ In cancer cell lines, constitutive Ca^2+^ flow from ER to mitochondria regulates OXPHOS activity.^40^ In permeabilized cells, we mimicked cytosolic [Ca^2+^] ranging from 0 nM to 8.4 µM. Interestingly, MICU2 was involved in the proper functioning of the respiratory chain and the production of ATP in the absence of cytosolic Ca^2+^ or at a physiological concentration of 150 nM. Surprisingly, we demonstrated that loss of MICU2 expression results in a specific defect in the activity and expression of complex IV (cytochrome *c* oxidase), limiting supercomplex assembly and finally maximal OXPHOS capacity, as determined on permeabilized cells, even in absence of cytosolic Ca^2+^. Paupe et al.^41^ also reported the link between mitoCa^2+^ uptake and defective complex IV assembly. The suppression of CCDC90A, also named MCUR1, in human fibroblasts produces a specific cytochrome *c* oxidase assembly defect, resulting in decreased mitoCa^2+^ uptake capacity. The consequences of MICU2 KO on ATP production in Ca^2+^-free conditions, and its effect on complex IV activity, may also point to a role for MICU2 in the respiratory chain, independent of Ca^2+^ and possibly independent of MCU. Additional experiments are required to determine this possible role.

### The pivotal role of MICU2 in the metabolic orientation

Mitochondria are the main mediators for the change in metabolic activity of cancer cells.^42^ This metabolic reprogramming is implicated in cancer progression and therapy resistance.^43^

CRC mostly arises from progressive accumulation of somatic mutations within cells. APC, KRAS, TP53, and MYC have been reported to participate in genetically global metabolic reprogramming.^44^ Pyruvate, produce through glycolysis, glutamine, and fatty acid metabolism form the substrates of the TCA cycle. All these pathways are altered in CRC.^45^ Glucose deprivation contributes to the development of KRAS pathway mutations in tumor cells.^46^ Oncogenic KRAS decouples glucose and glutamine metabolism to support cancer cell growth.^47,48^ p53 regulates biosynthesis through direct inactivation of glucose-6-phosphate dehydrogenase.^49^ MitoCa^2+^ uptake is essential to activate Ca^2+^-dependent dehydrogenases of the TCA cycle and to increase OXPHOS. In CRC cells, MCU expression promotes cell proliferation *in vitro* and *in vivo* by upregulating mitoCa^2+^ uptake and energy metabolism.^50^ .Using pharmacological blockers of the glutamine pathway or cell culture media devoid of glutamine, we observed that this substrate is essential for the viability of CRC cells. Furthermore, the oxidative metabolism and mitochondrial ATP production sustain through glutaminolysis is reduced by the down-expression of MICU2.

Here, we also described increased lipid accumulation in MICU2 KO cells and a reduction in the use of beta-oxidation related to proliferation mechanisms. Wang et al.^51^ showed that CPT1A-mediated activation of fatty acid oxidation increases the metastatic capacity. We also reported that blocking CPT1 and beta-oxidation with etomoxir and trimetazidine reduced the viability of Control KO cells but did not affect the viability of MICU2 KO cells. Recently, Tomar et al.^52^ demonstrated that MCU and mitoCa^2+^ shape bioenergetics and lipid homeostasis in hepatic cells.

Glucose is the primary source of energy of the cells and it supports important metabolic intermediates, and this contribution is thought to exert great effects on tumor cell metabolism. A decrease in mitochondrial ATP production implies an increase in energy synthesis by glycolysis. Because of their very different efficiencies, this means an increase in glucose consumption. In this study, we showed an increase in glucose uptake in MICU2 KO CRC cells. Furthermore, 2DG treatment led to a greater reduction in the viability of MICU2 KO cells than Control KO cells. These results corroborate the increase in lactate production in MICU2 KO cells, suggesting more glycolytic metabolism in MICU2 KO cells. Interestingly, inhibition of MPC with UK5099 did not affect the viability of MICU2 KO cells. Thus, in the absence of MICU2, cells develop a preference for glycolysis as the main energy supply. MitoCa^2+^ uptake is essential to activate Ca^2+^-dependent dehydrogenases of the TCA cycle and to increase OXPHOS. We demonstrated that MICU2 KO decreased mitoCa^2+^ influx in CRC cells; this modification inhibited the phosphorylation of PDH and the amount of acetyl-CoA.

The role of different MICU isoforms on bioenergetic preference is not very clear. Chakraborty et al.^31^ recently demonstrated that MICU1 overexpression in ovarian cancer cells induces both glycolysis and chemoresistance and that MICU1 overexpression in patients with ovarian cancer correlates with poor overall survival. In addition, MICU1 negatively regulates OXPHOS function. In the vast majority of cases, cancer cells undergo metabolic reprogramming and preferentially use glucose via aerobic glycolysis.^53^ In contrast, we have defined a role of MICU2 in maintaining an OXPHOS phenotype and the respiratory complex assembly in CRC cells at an advanced stage. Taken together, we propose that association between MICU1/MICU1, MICU2/MICU1, and MICU3/MICU1 implicated in MCU regulation shape the metabolic orientation of cancer cells. Thus, high MICU1 expression is associated with a glycolytic phenotype, while high expression of MICU2 favors respiration and mitochondrial metabolism. However, an MCU-independent contribution of MICU2 cannot be ruled out. Indeed, a recent report described MICU1 as an intermembrane space Ca^2+^ sensor that modulates mitochondrial membrane dynamics independently of matrix Ca^2+^ uptake.^54^ Therefore, it is possible that MICU exerts a similar role in specific circumstances.

In summary, we have demonstrated a specific role for the MICU2 in the regulation of mitoCa^2+^ uptake in CRC. We have also defined a crucial role for MICU2 in maintaining the integrity of the mitochondrial network and in the proper assembly and functioning of the respiratory chain. For the first time, we have demonstrated the central role of MICU2 in the balance between glycolysis and OXPHOS during cancer progression, and more specifically in CRC. This work provides a promising perspective to better understand and target metabolic diseases including cancers. The MICU2/MICU1 ratio could represent a singular predictive marker of the metabolic preferences and pave the way for the development of personalized metabolic therapies.

## Acknowledgments

The authors acknowledge H2P2 platform for technical assistance for immunohistochemistry analysis (Nicolas Mouchet, Anthony Sébillot, Gevorg Ghukasyan – Université Rennes 1) and PHENOMIN-TAAM-UPS44, CIPA (Alain Le Pape and Sharuja Natkunarajah - Centre d’Imagerie du Petit Animal, part of MOV2ING platform, Orléans) for *in vivo* models. Alison Robert is a recipient of a 3-years doctoral grant from Inserm / Région Centre Val-de-Loire. David Crottès is a recipient of Smart Loire Valley fellowship from Le Studium Loire Valley Institute for Advanced Studies. Thanks to Frederic Picou, member of the LNOx and Jacques Dupuy, INRAe Toxalim, for their help. Thanks to Agathe Brugoux, Oceane Pertegaz, Beatrice Genova, Cyrille Guimaraes and Michelle Pinault members of the N2C, for their technical help.

This project was partly supported by grants on behalf of the following french department committees of Ligue Contre le Cancer "Grand-Ouest": 16 (Charente), 37 (Indre-et-Loire), 49 (Maine-et-Loire), 72 (Sarthe) and 85 (Vendée). We acknowledge Inserm, INCa, Université de Tours and Région Centre-Val de Loire for their financial supports.

Inserm UMR 1069 is leader of Cancéropôle Grand-Ouest 3MC network (Marine Molecules, Metabolism and Cancer), member of Région Centre – Val de Loire thematic research consortium RTR MOTIVHEALTH (Molecular and Technological Innovation for Health) and member of CNRS research group APPICOM (GDR2082 - Integrative Approach for a multiscale functional analysis of membrane proteins). Alison Robert, Maxime Guéguinou, Arnaud Chevrollier, Naig Gueguen, Jean-François Dumas and William Raoul are also members of Meetochondrie French network.

## Ethics

Animal experimentation: This study was performed in strict accordance with the recommendations in the Guide for the Care and Use of Laboratory Animals. All of the animals were handled according to approved institutional animal care and use committee (authorisation Apafis 19933, regional ethical committee CEEA – 003 CNRS, Orleans) protocol. Every effort was made to minimize animal suffering.

## Author contributions

M.G. designed the project. A.R, D.C. J.B, I.D, A.C, N.G., M.G. performed cell culture experiments. A.R. J.B., S.S.,J.F.D., C.G., M.P. performed metabolic experiments. D.C, S.C performed bioinformatics analyses. A.R., D.C., W.R. M.G. wrote the manuscript, and all authors contributed to revision and edits. Funding Acquisition: T.L., C.V., W.R., M.G..

## Methods

### Transcriptomic analysis

Transcriptomic analyses and heatmaps were generated using R software. RNA-Seq data from colon cancer and normal colon were generated by The Cancer Genome Atlas Research Network (http://cancergenome.nih.gov/). Read counts and clinical annotation were download from GDC using the *tcgaworkflow* package. Read counts were then filtered, normalized, processed, and logarithmically transformed using the *edgeR* package. Primary tumors of TCGA-COAD were classified as “High” or “Low” if the value of their expression is respectively superior to the 95^th^ percentile or inferior to the 5^th^ percentile of their expression in normal samples. Heatmaps were generated using the *ComplexHeatmap* package. Differential expression analysis was performed on read counts using the *edgeR* package. DEA was obtained by comparing groups defined according to the expression of MICU2, MICU1, or the MICU2/MICU1 ratio. GSEA was performed on the log_2_(fold change)-ranked gene lists obtained by DEA using the *fgsea* package and gene set collections from MSigDB (https://www.gsea-msigdb.org/gsea/msigdb-c5.go.bp.v7.5.1.symbols.gmt). Published transcriptomic data of normal, primary, and metastatic CRC generated by microarrays (GSE41258) were used^55^. Raw data were downloaded and RMA pre-processed using the *affy* package.

### Cell culture

The CRC cell lines HT29, DLD1, HCT116, SW480, and SW620 were chosen to investigate the role of MICU2. CRC cells were cultured in RPMI-1640 media + GlutaMax (Gibco, Thermo Fisher, Illkirch, France) supplemented with 10% fetal bovine serum (FBS; Eurobio, Les Ulis, France) and 1% penicillin and streptomycin (Eurobio). NCM356 cells (Incell Corporation, LLC, San Antonio, TX, USA) were cultured in high-glucose Dulbecco’s Modifed Eagle Medium (DMEM) (Sigma-Aldrich, Missouri, USA) supplemented with 10% FBS and antibiotics. All cells were cultured in an incubator at 37°C with 5% CO_2_.

### Generation MICU2 KO

Several MICU2 KO clones in HCT116-Luc cells were generated by Ubigene (China) using the CRISPR/Cas9 system. MICU2 KO was generated using two exon 2 targeting sequences, MICU2-gRNA3 (ACACTTAGAGATTAAACGAGG) and MICU2 gRNA6 (GTATTCCAGTACACTTAGGAAGG). All clones were validated by polymerase chain reaction (PCR) and imaging.

### Metabolic pathway inhibitors

The following inhibitor references are described in the key resources table: 2DG (5mM), 2NBDG (100µM), UK5099 (50µM), rotenone (0.5, 1, and 2.5µM), oligomycin (0.5, 1, and 2.5µM), trimetazidine (10µM), etomoxir (10µM), telaglenastat (1 and 5µM), V-9302 (5 and 10µM), BAPTA (2.5 and 5µM), and MITOTEMPO (2.5 and 5µM).

### Specific media

Specific media were used: RPMI Medium 1640 [-] Glutamine (Reference: 42401018, Gibco, Thermo Fisher, Illkirch, France), RPMI Medium 1640 [-] Glucose (Reference: 11879020, Gibco, Thermo Fisher, Illkirch, France), RPMI Medium 1640 [+] 4.5 g/L D-Glucose (Reference: A1049101, Gibco, Thermo Fisher, Illkirch, France).

### Cell proliferation

After harvesting, 3 × 10^3^–5 × 10^3^ cells were plated in each well of 96-well plates. The platers were incubated at 37°C in 5% CO2 for 4 h to allow the cells to adhere to the plate. The CyQUANT-NF dye was diluted in HBSS buffer, and 100 µL of the mixture was added in each well. The plate was kept at 37°C in 5% CO2. The fluorescence intensity (∼485/∼530nm) was measured using FlexStation 3 Multimode Plate Reader (Molecular Devices, California, USA).

### Cell viability

Cell viability, survival in HBSS, and short-term toxicity were evaluated using standard sulforhodamine B (SRB) method after treatment for 24 and 48h. Briefly, cells were fixed with 50% trichloroacetic acid for 1h at 4°C and stained for 15min with 0.4% SRB solution. Cells were then washed three times with 1% acetic acid and dye was dissolved using 10mM Tris over 10min. Absorbance at 540nm was determined with a BioTek (Vermont, United States) Spectrophotometer.

### Cell cycle

Cells were harvested using trypsin-ethylenediaminetetraacetic acid (EDTA, Thermo Fisher Scientific, San Jose, CA, USA) 24h after treatment, washed in 1× phosphate-buffered saline (PBS), fixed in cold 70% ethanol, and incubated at -20°C for at least 2h. Subsequently, the samples were resuspended in PBS with RNase and stained with 0.025 mg/mL propidium iodide (PI) for 30min in the dark at room temperature. The DNA content of stained cells was analyzed using a Gallios flow cytometer (Beckman Coulter, Villepinte, France). For each sample, a minimum of 5×10^4^ cells were evaluated. Analyses were done using the Kaluza 1.3 software (Beckman Coulter).

### Cell migration in the wound healing assay

First, 6×10^5^ cells were seeded on each side of an Ibidi culture insert (Ibidi GmbH, Gräfelfing, Germany) in 100µL of RPMI supplemented with 10% FBS and enriched with 1% antibiotics. The 24-well plate was incubated at 37°C and 5% CO_2_ for 24h. The insert was then removed to create the scar and 500µL of medium was added. The plate containing the scarred cells was then placed in a chamber maintained at 37°C and 5% CO_2_, connected to a camera so that photographs of the scar could be taken at regular intervals for 24h. At the end of the migration, the area of the scar not covered by cells was measured using the T-scratch software. The percentage of coverage is calculated using the formula:

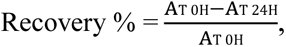

where A_T 0H_ is the uncovered scar area measured at time T0 immediately after scar creation and A_T 24H_ is the uncovered scar area measured at time T24, 24h after scar creation.

### Spheroid base migration

Spheroids were prepared by seeding drops of cells at a rate of 3×10^3^ cells per 30µL into the lid of a petri dish (Reference: 353003, Falcon), the lid was then turned over to form a spheroid. After 4 days, the spheroid was placed in a 96-well plate (Reference: 353077, Falcon) coated with fibronectin and the migration around the spheroid was photographed using a microscope (Nikon, Champigny-sur-Marne, France) at 0, 24, and 48h.

### Subcutaneous CRC xenograft tumor model

CRC xenografts were established by injecting 5×10^6^ cells per Swiss nude mouse (n=10 per cell line) as described previously by Gueguinou et al.^56^ All mice were assessed weekly using whole-body bioluminescence imaging and photoacoustic imaging as described below. Authorization (Apafis 19933) was given by the regional ethical committee CEEA – 003 (Campus CNRS, Orléans) and the French Ministry of Higher Education, Research and Innovation (MESRI).

### Ultrasound/photoacoustic imaging, bioluminescence imaging, and hypoxia immunostaining

The protocols have already been described in detail.^57^ Briefly, bioluminescence imaging was performed once a week until the end of the study using an IVIS-Lumina II (Perkin Elmer, France). Each mouse was injected intraperitoneally with 100mg/kg luciferin potassium salt (Promega, France). After mice were anesthetized, acquisition binning and duration were set depending on tumor activity. Signal intensity was quantified as the total flux (photons/seconds) within regions of interest drawn manually around the tumor area using Living Image 4.4 software (Perkin Elmer).

For photoacoustic imaging, mice were anesthetized with 1.5% isoflurane and placed on a thermostatically heating pad (37°C). Respiratory gating was derived from electrocardiography. A colorless aqueous warmed ultrasonic gel (Supragel1, LCH, France) without any air bubbles was applied between the skin and the transducer. Tumors were imaged with the VisualSonics VevoLAZR System (FUJIFILM VisualSonics Inc, Canada). Three-dimensional (3D) scans were made of digitally recorded ultrasound images. The tumor area was measured by delineating the margins using Vevo1LAB 3.2.6 software. For hypoxia assessments, tumors were investigated by photoacoustic imaging with the OxyHemo-Mode to determine the average SO2 values and the corresponding hypoxic volumes. A transducer with central frequency at 21 MHz was used for B-Mode imaging and photoacoustic imaging. Pimonidazole immunostaining was performed with the Hypoxyprobe kit (Hypoxyprobe, Burlington, USA), following the supplier’s instructions. Briefly, mice received an intravenous injection of 60 mg/kg of the pimonidazole solution and were sacrificed 60 min following the injection. The tumors were then resected and fixed for 24 h in 10% formaldehyde. The tumors were embedded in paraffin and sectioned. The primary antibody was mouse anti-pimonidazole and the secondary antibody was FITC-labeled goat anti-mouse (Abcam, Cambridge, USA).

### Lipid droplet staining

Forty-eight hours before fixation, 1 × 10^5^ cells were placed per well in an 8-well cell culture chamber (Reference: 154534, Labtek, Brendale, QLD). The cells were fixed with a 4% paraformaldehyde at room temperature for 15 min. Cell culture chambers were rinsed twice with cold PBS, permeabilized with 0.5% Triton X-100 for 5 min at room temperature, rinsed three times with PBS, and then incubated with blocking solution (PBS containing 0.2% bovine serum albumin [BSA], 0.02% sodium azide, 0.05% Triton X-100, and 10% FBS) for 1 h at room temperature. Cells were stained with 0.3% Oil Red O (diluted in 60% isopropanol) for 2 min at room temperature. The cell culture chambers were then washed twice in PBS and nuclei were stained with DAPI for 5 min. The cell culture chambers were then washed once more and mounted with coverslips using SlowFade Gold antifade mounting medium (Reference: P36961, Molecular Probes, Oregon, USA).

Oil Red O was used to quantify the number of lipid droplets. The images were acquired with a Leica SP8 microscope (microscopy department of the University of Tours). Image processing and analysis were performed using ImageJ/Fiji software (National Institutes of Health, Bethesda, MD, USA). After subtracting the background (using a 20-pixel rolling ball radius) and setting an identical threshold for all images, the particle analysis function was used to measure the number of lipid droplets per cell.

### Immunohistochemistry

Immunohistochemistry was done with a Discovery Ultra Machine (Roche, Mannheim, Germany) and the Chromo-Map Kit (Reference: 760-159, Roche). The primary antibody (Ki67) was diluted with Antibody Diluent (Reference: 760-108, Roche) and the samples were incubated 60 min at 37°C. Automatic application of secondary antibody was done with Omni Map anti-Rb HRP (Reference: 760-4311, Roche). The staining was visualized by automatic addition of the substrate (H_2_O_2_+DAB) of the Chromo-Map Kit and the HRP of the secondary antibody. Counter-staining was performed with Hematoxylin II (Reference: 790-2208, Roche) for 16 min and Bluing Reagent (Reference: 760-2037, Roche) for 4 min. The slides were then dehydrated and mounted with Pertex mounting media (Reference: 00811-EX, Histolab, Askim).

### 3D fluorescence microscopy

Cells were transfected with a mitochondria-targeted fluorescent protein (PDH_GFP) or incubated for 15 min with 100 nM Mitotracker green (Molecular Probes) to stain the mitochondrial network. For fluorescence imaging, coverslips were mounted in housing and placed on the stage of an inverted wide-field ECLIPSE Ti-E microscope (Nikon, Tokyo, Japan) equipped with a 100× oil immersion objective lens (Nikon Plan Apo100x, N.A. 1.45) and an Andor NEO sCOMS camera controlled by Metamorph® 7.7 software (Molecular Devices, Sunnyvale, CA, USA). A precision piezoelectric driver mounted underneath the objective lens allowed faster Z-step movements, keeping the sample immobile while shifting the objective lens. Fifty-five image planes were acquired along the Z-axis at 0.1-μm increments. For mitochondrial network characterization, the acquired images were iteratively deconvolved using Huygens Essential® software (Scientific Volume Imaging, Hilversum, the Netherlands), with a maximum iteration score of 50 and a quality threshold of 0.01. Imaris 8.0® software (Bitplane, Zurich, Switzerland) was used for 3D processing and morphometric analysis. The mitochondrial network was modelled in 3D, and thresholds were defined to classify mitochondria depending on their volume (Imaris Isosurface Tools).

### Reverse transcription real-time polymerase chain reaction (RT-qPCR)

Total RNA was collected using the Nucleospin RNA Kit (Macherey–Nagel, Hoerdt, France) and transcribed into complementary DNA (cDNA) with the PrimeScript RT Reagent Kit (RR037A, Takara, Kusatsu, Japan). cDNA was then amplified with the SYBR Green Master kit (Roche) using a Light Cycler 480 apparatus. RT-qPCR was performed in 40 cycles of 95°C for 15 s and 60°C for 45 s. The average ΔCt value was calculated for each cell line with respect to the housekeeping gene HPRT1. The primers sequences used are:

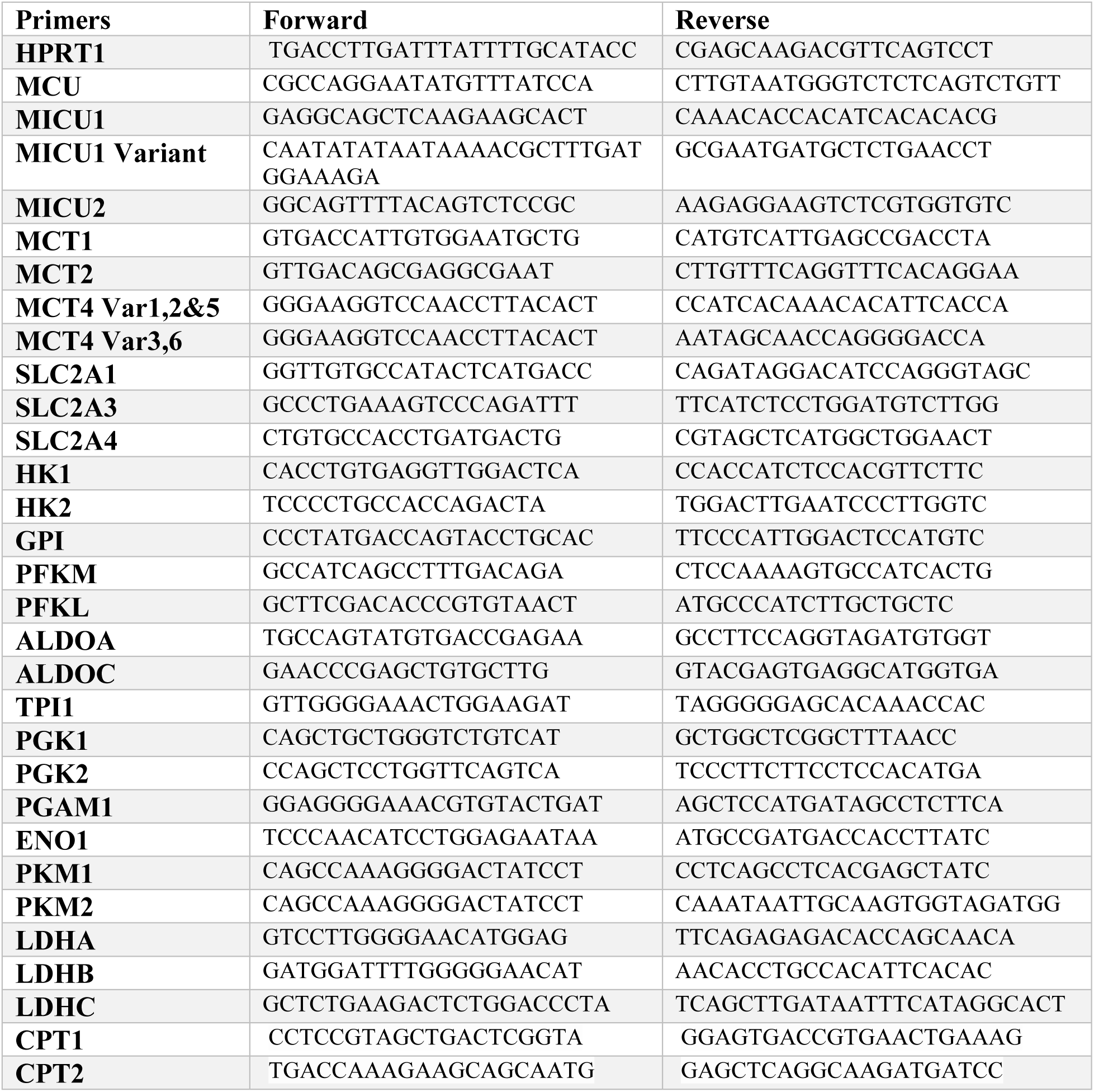

### Western blot analysis

HT29, DLD1, SW480, HCT116, and SW620 colon cancer cells and NCM356 non-tumoral cells were collected with a lysis buffer containing protease inhibitors and phosphatase inhibitors and used for protein assay. A BCA protein assay kit (Reference: 23227, Thermo Fisher Scientific, France) was used to determine the protein concentration. The resulting proteins were separated by polyacrylamide gel electrophoresis (4%–15% Mini-PROTEAN® TGX Stain-Free™ Protein Gels, Bio-Rad, California, USA) and then transferred to a nitrocellulose membrane (Reference: 1704158, Bio-Rad). The membranes were incubated in primary antibody diluted 1:1000 in TBST with 5% milk overnight at 4°C with agitation. The primary antibodies were against MCU (Sigma-Aldrich), MICU2 (Abcam), MICU1 (Sigma-Aldrich), MFN2 (Cell Signaling, USA), OPA1 (BD Biosciences, France), DRP1 (Cell Signaling), PDRP1 (Cell Signaling). The following day, the membranes were washed three times with TBST and incubated with HRP-conjugated secondary antibody at room temperature for 1 h with shaking. The secondary antibodies used were: m-IgG Fc BP-HRP (1:5000, sc-525409) and Ms x Rb Light Chain specific HRP (1:2000, MAB201P). The internal control was total protein deposited and quantified with a gel stain. An ECL chemiluminescence kit (Clarity Western ECL substrate, Bio-Rad) was used to visualize the bands, which were quantified with Image Lab software (Bio-Rad).

### MitoCa^2+^ measurements

To measure mitoCa^2+^ using mito-SP-linker-GCaMP6m, named mt-riG6m, cultured cells were transfected using Lipofectamine 2000 (Invitrogen, USA) with 1.5 µg of pCDNA3.1 mt-riG6m 2 days before imaging. Twenty-four hours before the experiments, transfected cells were plated in FluoroDish™ FD35-100. The cells were stimulated with 10 µM LPA in PBS containing 2 mM CaCl_2_. Time-lapse images were acquired using epifluorescence microscopy (Nikon). mt-riG6m and miGer were gifts from Li et al.^58^

ER Ca^2+^ store depletion was quantified by transfecting parental and MICU2-KO HCT116 cells with red miGer^58^ using Lipofectamine 48 h prior to imaging. Cells expressing miGer were then excited at 552 nm and relative ER Ca^2+^ measurements were recorded through a 20× objective lens. Immediately after identifying R-miGer-positive cells, the bath solution was replaced with nominally Ca^2+^-free HBSS. One minute into the experiment, the cells were treated with TG (Reference: T7458, Life Technologies, Thermo Fisher).

### SOCE measurement by Fura-2 AM

Intracellular Ca^2+^ imaging was performed as described previously.^59,60^ Cells were plated in 96-well plates at 2 × 10^4^ cells per well 24 h before the experiment. Adherent cells were for loaded with the ratiometric dye Fura2-acetoxymethyl ester (AM; 2 μM) at 37°C for 45 min and then washed with PBS supplemented with Ca^2+^. During the experiment, the cells were incubated with Ca^2+^-free physiological saline solution (PSS) solution and treated with 2 µM TG to deplete intracellular store of Ca^2+^. Ca^2+^ entry was stimulated by injecting 2 mM of CaCl_2_. Fluorescence emission was measured at 510 nm using the FlexStation-3 (Molecular Devices) with excitation at 340 and 380 nm. The maximum fluorescence (peak of Ca^2+^ influx [F340/F380]) was measured in MICU2 KO cells and compared with the Control cells.

### Measurement of oxygen consumption

#### Seahorse analysis

The cellular oxygen consumption rate (OCR) data were obtained using a Seahorse™ XF96 Flux Analyzer from Seahorse Bioscience (Agilent Technologies, Santa Clara, CA, USA). The experiments were performed according to the manufacturer’s instructions. Briefly, HCT116 cells were seeded in XF96 cell culture plates at 2×10^4^ cells/well. On the day of analysis, the culture medium was replaced with XF DMEM (Thermo Fisher Scientific, San Jose, CA, USA) supplemented with 2mM glutamine and lacking bicarbonate (pH 7.4). The cells were then incubated at 37°C in a non-CO_2_ incubator for 1h. Measurements were made as described in the relevant figure legends. Sequential injection of 10mM glucose, 1µM oligomycin, 100µM dinitrophenol (DNP), and 0.5µM rotenone/antimycin A permitted the determination of the main respiratory parameters. Finally, the data were normalized to the amount of DNA present in the cells and assayed using the Cyquant® Cell Proliferation Assay kit (Thermo Fisher Scientific, San Jose, CA, USA). The data were acquired with the Seahorse Wave Controller and analyzed with the Seahorse Wave Desktop Software.

#### Whole cell measurement

The measurements were performed at 37°C under magnetic stirring by placing 2×10^6^ cells in a final volume of 2mL of culture medium (RPMI, RPMI without glucose, or RPMI with high glucose) with 10% FBS in an oxygraph (OROBOROS oxygraph-2k). The cells of each line were treated with their respective medium 48h before the experiment. They were then harvested, counted, and centrifuged at 700*g* for 3min. The cell pellet was suspended in 5 mL of selected medium with FBS and centrifuged at 800*g* for 2min. The resulting residual pellet was then resuspended in 500μL of selected medium with SVF (cell solution used for the oxygenation measurement). The resuspended cells were added to 1.5mL of different medium with FBS already present in the measuring tank. As soon as a steady slope was obtained, the oxygen consumption was measured. OXPHOS was then inhibited by adding oligomycin (an ATP synthase inhibitor; 5.25μg/mL); the respiration rate obtained corresponds the energy wasting state. Successive additions of FCCP (a protonophoric uncoupler; 0.8μM) were then made until the maximum respiration rate was reached. Finally, 2µM antimycin A was added to determine non-mitochondrial oxygen consumption.

#### Measurement of permeabilized cells

Measurements were performed at 37°C—with agitation, in a final volume of 2 mL of respiration buffer with BSA and different [Ca^2+^] (0, 500 nM, 1 µM and 8.4 µM) with 3× 10^6^ cells (OROBOROS oxygraph-2k) previously permeabilized with digitonin. The cells of each line were detached with trypsin, counted and centrifuged at 700*g* for 3min. The pellet was suspended in 1mL of ASB-free respiration buffer and the plasma membrane was permeabilized by adding digitonin (1mg/mL previously heated at 95°C for 2.5min) (6μL per 1 × 10^6^ HCT116 cells). The whole set was gently shaken for 2 min. Then, 4 mL of respiration buffer (10 mM KH_2_PO_4_, 300mM mannitol, 10mM KCl, 5mM MgCl_2_, and 1mM EGTA, pH 7.4 at 37°C) with ASB fatty acid free was added to stop the action of digitonin. The mixture was centrifuged at 800*g* for 2min. The resulting pellet was then resuspended in 500μL of respiration buffer with BSA (solution for oxygen measurement). The cells resuspended in the respiration buffer with BSA were added to the measuring cell containing 1.5mL of respiration buffer with BSA. As soon as a steady slope was obtained, succinate (10mM) and ADP (1.5mM) were added and the oxygen consumption related to ATP synthesis was determined. OXPHOS was then inhibited by adding oligomycin (5.25μg/mL). Once the oxygen consumption stabilized, FCCP (0.8µM) was added until the maximum respiration was reached. Cytochrome *c* was added, to control the integrity of the outer mitochondrial membrane and then 2µM antimycin A was added to inhibit complex III.

### Expression of super complexes

For supercomplex assembly analyses, mitochondria were enriched by using differential centrifugation, according to a slightly modified version of the method described by Bonnet et al.^61^ Briefly, the cell pellets were incubated for 10 min on ice with cold digitonin (4mg/mL in PBS [v/v] or 0.2% w/v) to dissolve cell membranes. Then, digitonin was diluted by adding cold PBS (5% v/v). Cells were centrifugated at 10,000*g* for 10min at 4°C to recover the mitochondria-enriched fraction. The pellet was washed once more in 1mL of cold PBS, centrifuged (10,000*g* at 4°C), resuspended at 10 mg/mL in AC/BT buffer (1.5M aminocaproic acid and 75mM Bis-Tris/HCl, pH 7.0, supplemented with Complete Mini Protease Inhibitor [Roche Diagnostics, Stockholm, Sweden]), and kept frozen at -80°C until analysis. For blue native polyacrylamide gel electrophoresis (BN-PAGE) analyses, 50µg of each sample was diluted at 2mg/mL in AC/BT buffer and mitochondrial membrane proteins were solubilized by incubation with 6g/g protein digitonin (1.2% w/v) for 10min at 4°C. After centrifugation at 20,000*g* for 20min at 4°C, the supernatant was collected, and 5% Serva Blue G dye (Bio-Rad) in 1M aminocaproic acid/50mM Bis–Tris/HCl, pH7.0 was added (1/20 v/v) prior to loading. Respiratory chain complexes and supercomplexes were separated on Native PAGE Novex 3%–12% Bis-Tris gels (Invitrogen) for approximatively 3 h and transferred to polyvinylidene fluoride membranes (GE Healthcare, Velizy-Villacoublay, France) in cold blotting buffer (25mM Tris, 192mM glycine, pH 8.3, and 20% methanol). Membranes were hybridized using dedicated monoclonal antibodies (NDUFS2 rabbit monoclonal, SDHA rabbit monoclonal, UQCRC2 mouse monoclonal, MT-CO1 (COX1) mouse monoclonal, Abcam, the Netherlands). Acquisitions were performed using the Odyssey FC imaging system and analysed using Image Studio™ Lite (LI-COR Biosciences).

### Mitochondrial enzymatic activities

The activities of complex I (NADH ubiquinone reductase) and citrate synthase (CS) were measured at 37°C with a UVmc2 spectrophotometer (SAFAS, Monaco) in a mitochondria-enriched fraction. Cells were resuspended in cell buffer (250mM saccharose, 20mM tris[hydroxymethyl]aminomethane, 2mM EGTA, 1mg/mL BSA, pH 7.2; 1×10^6^ cells per 50µL), disrupted by two freeze-thaw cycles, washed, and centrifuged at 16000*g* for 1min to eliminate the cytosolic fraction. For complex I and V analyses, cells were further resuspended in the cell buffer (1×10^6^ cells per 250µL) and sonicated (six times × 5s) on ice. The activities of respiratory chain complex activities were measured according to standard routine clinical protocols for CI (NADH:ubiquinone reductase, NUR),^62^ complex II (succinate:ubiquinone reductase, SUR), Complex III (ubiquinol:cytochrome *c* reductase, UCCR), Complex IV (cytochrome *c* oxidase, COX)^63^, Complex V (F1-ATPase), and CS^63^. The protein content was determined with the bicinchoninic assay kit (Uptima, Interchim, Montluçon, France) using BSA as the standard.

### PDH activity assays

The PDH activity of MICU2 KO cells was measured with the colorimetric PDH Assay Kit (Sigma-Aldrich) according to the manufacturer’s instructions.

### Acetyl-CoA ratio measurement assay

The acetyl-CoA content of the cell culture medium was determined with the Acetyl-CoA Assay Kit (Reference: ab87546, Abcam) according to the manufacturer’s instructions.

### Lactate production

Lactate production was assessed with the L-Lactic Acid Colorimetric Assay Kit (Reference: E-BC-K044-S, Elabscience, Texas, USA) according to the manufacturer’s instructions.

### Fatty acid oxidation activity

Visualization of fatty acid oxidation in the HCT116 Control and KO MICU2 cells lines treated with FAOBlue (Funakoshi, Japan) in HEPES-buffered saline (HBS) buffer with or without pre-treatment with etomoxir (20μM, 3h), a potent fatty acid oxidation inhibitor. After FAOBlue incubation, blue fluorescence (excitation 405nm, emission 430–480nm) was observed.

### Detection of ROS

A total of 1 × 10^6^ cells were stained for 10min at 37°C with 5 µM DCFDA and then washed and analysed by flow cytometry (excitation ∼492–495nm, emission 517–527nm) to detect cytosolic ROS. Mitochondria-derived ROS were detected in 1×10^6^ cells stained at 37°C for 10min with 5µM MitoSOX (DHE). After staining, the cells were washed and analysed by flow cytometry (excitation 510nm, emission 580nm). The cells were analysed immediately after completing staining.

### Statistical analysis

The data are presented as the mean ± standard error of the mean and were analyzed with GraphPad Prism 6 (GraphPad Software, San Diego, CA, USA). Statistical analysis was performed with a paired Mann–Whitney test, the Kruskal–Wallis test, or analysis of variance (ANOVA) followed by Dunnett’s, Dunn’s, or Sidak’s multiple comparisons test. The exact tests are indicated in the figure legends. A p-value <0.05 was considered significant.

