## Supplemental data for "Encoding of the colorectal cancer metabolic program through MICU2"

**Figure S1. MICU2 KO and expression of MCU, MICU1 and MICU2/MICU1 ratio in CRC cells**
(A) Transcriptomic analysis of the MICU2/MICU1 ratio according to CRC stages in the TCGA-COAD (B) Transcriptomic analysis of the MICU1 and MICU2/MICU1 ratio in the GSE41258 datasets. Each data point represents an individual sample (ANOVA followed by Dunn's multiple comparisons test). (C) Bar plot representing the expression of MCU, MICU1, MICU1 variant, and MICU2 mRNAs measured by RT-qPCR in the HCT116 Control and MICU2 KO cell lines (n=3, ANOVA followed Dunnett's multiple comparisons test, **p<0.01 and ***p<0.001). (D) Density plots representing the definition of the status of MICU1, MICU2, and the MICU2/MICU1 ratio of primary colon tumor samples of the TCGA-COAD dataset.

**Figure S2. The role of MICU2 in cell proliferation, migration, and sensitivity to chemotherapy**
(A) Heatmap of the most differentially expressed genes between colon tumor samples in the TCGA-COAD dataset with low, normal, or high expression of MICU2/MICU1 ratio. (B) Heatmap of normalized enrichment scores of all cell cycle–associated genes obtained by GSEA for primary colon tumor samples with low, normal, or high expression of MICU2. A star indicates an adjusted p-value <0.05. (C) Boxplots representing the viability of the Control and MICU2 KO cell lines cultured in the absence of presence in 5FU (500nM, 1µM, 5µM, and 10µM) (top panel) and oxaliplatin (500nM, 1µM, 5µM, 10µM, and 20µM) (bottom panel) for 24 and 48h (n=9–11). (D) Left panel: representative images of the wound healing assay. Right panel: curve and boxplots representing the percentage of gap closure as a function of time and after 24h in the Control and MICU2 KO cell lines. The scale bar is 5µm (n=6-8, Kruskal–Wallis test). (E) Left panel: representative images of 3D spheroids of the Control and MICU2 KO cell lines plated on top of a fibronectin layer after 0, 24, and 48h. Right panel: boxplot representing the migration area at 24 and 48h. The scale bar is 20µM (n=5-8, Kruskal–Wallis test). On all plots, *p<0.05, **p<0.01, and ***p<0.001.

**Figure S3. The effect of MICU2 knockout on mitochondrial dynamics and ROS/$Ca^{2+}$ signaling**
(A) Bar plots representing the fold change of the expression of genes associated with mitochondrial dynamics in the Control and MICU2 KO cell lines compared with the Control KO cell line (n=3–5). (SB) Left panel: representative measurements of cytosolic [$Ca^{2+}$] in the Control and MICU2 KO cell lines measured with Fura-2 AM in response to 1µM of LPA applied in the absence of extracellular $Ca^{2+}$ and in the presence of 2 mM of extracellular $Ca^{2+}$. Right panel: boxplot representing the cytosolic Fura-2AM 340/380 ratio measured in the presence of extracellular $Ca^{2+}$ in the Control and MICU2 KO cell lines (n=4, N=27–55). (C) Left panel: representative measurements of cytosolic [$Ca^{2+}$] in the Control and MICU2 KO cell lines measured with Fura-2AM in response to 2µM TG applied in the absence of extracellular $Ca^{2+}$ and in the presence of 2mM of extracellular $Ca^{2+}$. Right panel: boxplot representing the basal Fura-2AM 340/380 ratio measured in the presence of extracellular $Ca^{2+}$ in the Control and MICU2 KO cell lines (n=6, N=42–95, ANOVA followed by Dunnett's multiple comparisons test). (D) Bar plots representing the viability of the Control and MICU2 KO cell lines cultured 24 or 48 h in presence or absence of BAPTA-AM (a $Ca^{2+}$ chelator) (n=3–5, ANOVA followed by Dunnett's multiple comparisons test). (E) Boxplots representing ROS production in the Control and MICU2 KO cell lines. Total ROS, mitochondrial ROS, and hydrogen peroxide were measured using DHE, MitoSox, and DCFDA, respectively. (F) Bar plots representing the viability of the Control and MICU2 KO cell lines incubated for 24h with MITOTEMPO (a mitochondrial ROS chelator) (n=5–7, ANOVA followed by Dunnett's multiple comparisons test). On all plots, *p<0.05, **p<0.01, and ***p<0.001.

**Figure S4. Molecular and functional analysis of the mitochondrial respiratory chain**
(A) TCGA analysis showing expression values of respiratory chain complexes as a function of low, normal, and high expression of MICU1, MICU2, and the MICU2/MICU1 ratio. (B) Boxplot representing the average expression of genes associated with the assembly of mitochondrial respiratory chain complexes I, II, III, and IV based on the status MICU1 and MICU2/MICU1 ratio in primary colon tumor samples from the TCGA-COAD dataset. (C) The expression levels of MCU, MICU1, and MICU2 were determined in the Control KO and MICU2 KO cells treated with the inhibitors rotenone and oligomycin using RT-qPCR. The data are presented as the mean ± standard error of the mean of four independent experiments. On all plots, *p<0.05, **p<0.01, and ***p<0.001.

**Figure S5. The impact of MICU2 deletion on fatty acid metabolism and beta-oxidation activities**
(A) The expression of MCU, MICU1, and MICU2 were determined in the Control and MICU2 KO cell lines treated with the inhibitors etomoxir (left panel) and trimetazidine (right panel) using RT-qPCR. The data are presented as the mean ± standard error of the mean of three or four independent experiments. (B) Efficacy of V-9302 (ASCT2 inhibitor) (left panel) and telaglenastat (glutaminase inhibitor) (right panel) treatments (1–10µM) after 48h on the Control and MICU2 KO cell lines (n=5, ANOVA followed by Dunn's multiple comparisons test, $^{*}p<0.05$. (C) qRT-PCR data showing fold changes in mRNA levels of glycolysis regulator proteins in the Control and MICU2 KO cell lines (n=4–7, ANOVA followed by Dunnett's multiple comparisons test, $^{*}p<0.05$ and $^{**}p<0.01$). (D) RT-qPCR data showing fold changes in mRNA levels of pyruvate regulator proteins in the Control and MICU2 KO cell lines (n=3–5, ANOVA followed by Dunnett's multiple comparisons test, $^{**}p<0.01$). (E) RT-qPCR data showing fold changes in mRNA levels of pentose phosphates proteins in the Control KO and MICU2 KO cell lines (N=5, ANOVA followed by Dunnett's multiple comparisons test, $^{*}p<0.05$ and $^{***}p<0.001$).

**Figure S6. Expression of MICU2 related to genes involved in glycolysis, hypoxia, mitochondrial respiration, and the TCA cycle**
(A) The expression of MCU, MICU1, and MICU2 was determined in the Control and MICU2 KO cell lines treated with the inhibitor UK5099 using RT-qPCR. The data are presented as the mean ± standard error of the mean of four independent experiments. (B) Heatmap representing the expression of genes associated with glycolysis, hypoxia, mitochondrial respiration, and the TCA cycle as a function of the MICU2/MICU1 status of primary tumor samples from the TCGA-COAD dataset. (C) Boxplot representing the mean expression of genes associated with glycolysis, hypoxia, mitochondrial respiration, and the TCA cycle according to the MICU1 and MICU2/MICU1 ratio status of primary tumor samples of the TCGA-COAD dataset. (D) Scatter plot representing the correlation of the average expression of genes associated with glycolysis, hypoxia, mitochondrial respiration, and the TCA cycles as a function of the expression of MICU1, MICU2, or the MICU2/MICU1 ratio in a panel of 57 CRC cell lines from the RNA-Seq dataset of the CCLE. (R is the Pearson correlation coefficient). (E) Boxplots representing the average expression of genes associated with glycolysis, hypoxia, mitochondrial respiration, and the TCA cycle in a panel of 57 CRC cell lines from the RNA-Seq dataset of the CCLE classified as low or high expression of MICU1, MICU2, or the MICU2/MICU1 ratio as a function of the median expression of MICU1, MICU2 or the MICU2/MICU1 ratio. On all plots, $^{*}p<0.05$, $^{**}p<0.01$, and $^{***}p<0.001$.

**SUPPLEMENTAL METHODS**

**Reverse transcription and real-time polymerase chain reaction**

| Primers | Forward | Reverse |
|---|---|---|
| **MFN2** | TGAGAGGCATCAGTGAGGTG | GCAGAACTTTGTCCCAGAGC |
| **OPA1** | AATCGGACCCAAGAACAGTG | ACTCCTCGGGATTCAAGGTT |
| **FIS1** | GGAGGACCTGCTGAAGTTTG | ACGATGCCTTTACGGATGTC |
| **DRP1** | GGAACAGCGAGATTGTGAGG | CCTTTGGCACACTGTCTTGA |
| **PFKFB1** | ACATGGAAGCCCTGCAAAT | GGCTGAGATAGTTGTGAACATCC |
| **PFKFB2** | ATCTCTCGGGGTGCCCTAT | TGCATAGGTCATCTCTTCACACA |
| **PFKFB3** | CAACAGCTTTGAGGAGCATGT | GGGAGCCTTTCATGTTTTGT |
| **PFKFB4** | CCAGATGAAGAGGACAATCCA | TCCTCGTAGGTCATTTCCTCA |
| **PDK1** | CACCAAGACCTCGTGTTGAG | ACGTGATATGGGCAATCCAT |
| **PDK2** | CTGGCCAACATCATGAAAGA | CCAGGAGGCTCTGGACATAC |
| **PDK3** | TGTGTGAACAGTATTACCTGGTAGC | GTTTGTCTGGCGCTTTGG |
| **PDK4** | CAGTGCAATTGGTTAAAAGCTG | GGTCATCTGGGCTTTTCTCA |
| **PDP1 var 1** | ATGTTGTCGGCTCCGTGT | TGGAACTTCTGACTGGGATTC |
| **PDP1 var 5** | TCGGGAAGAATCGTTTGGT | AAAAACAGTTGAGTTGGTGCTG |
| **PDP2** | CTGAGCCTGAGGTCACATACC | AGGCCAGCACAAGGAACTTA |
| **PDPR** | GTGGCCTATCACCTCTCCAA | GCCAGCACAGAACCTGGTAG |
| **GCKR** | AGATGATATTCGGGCTGCTC | GGCGATGGGTATCACCTGT |
| **DLAT** | GGGTGAGAAGCTAAGTGAAGGA | CAGGGACCAGGATTTTTGC |
| **DLD** | CCCAGAGGTAGAATTCCAGTCA | GAGCCAGCATTGGACCAG |
| **PDHA1** | GTCCGAGAGGCAACAAGGT | AAGTCTGCAGCTCCATCAGG |
| **PDHB** | CGGATAGAGGACACGACCA | GTCCAGTGAAAGCGCCTCT |
| **DERA** | CTGCGGGCCATTAGAGATT | AGAGAGCCAAGCAAGGGAAT |
| **G6PD** | GCAAACAGAGTGAGCCCTTC | GGCCAGCCACATAGGAGTT |
| **H6PD** | GGGTGGAGATCATCATGAAAG | GCGAATGACACCGTACTCCT |
| **PGD** | ATTGCTGCAAAAGTGGGAAC | GTTGTGCACCATCTTCACGA |
| **PGLS** | GCTGAGGACTACGCCAAGAA | AGGATCAGCAGGTCGAAAAC |
| **PGM1** | GATGGACGCGAGCAAACT | GACAGCCCACAGTCCATCTT |
| **PRPS1** | GCATTGCTCCAAAATACAGGT | AAGGGACATGGCTGAATAGG |
| **PRPS2** | ACACTTGCGGCACCATCT | TCCCATGGGTAAGGATAGCA |
| **RBKS** | CTACAGGAACTGCTTCTATAATTGTCA | TCAGATCCTCCGTATTCAAAAGT |
| **RPE** | CGATGGTGGAGTAGGTCCTG | AGCACTGCCAGACACAATCA |
| **RPIA** | TCATCGTGATCGCTGATTTC | GATGACCTCGATGGGGATT |
| **TALDO1** | CAGATGCCCGCTTACCAG | TTTAATCTGGTCCTCTTGTGACC |
| **TKT** | GGATGACCAGGTGACCGTTA | CGCGGATGTTGATCTTTTCT |
| **UCP2 Blast 1** | AAGGGGATCGGGCCATGATA | GAGCTGGGTTGCAGGATGTT |
| **UCP2 Blast 2** | CAAAGCCGGCTGGGTCTTAT | ACTGAGAAGGCTCAGGCAAA |
| **UCP3 long** | AGCCCCCTCGACTGTATGAT | ACTTTCATCAGGGCCCGTTT |
| **UCP3 Short** | AGCCCCCTCGACTGTATGAT | AGGAGGCTCACCCCTTGTAG |
| **PRMT1 Cell** | CCAGTGGAGAAGGTGGACAT | GTCATTCCGCTTCACTTGCA |
| **PRMT1 Blast** | CGCGAACTGCATCATGGAGAA | CTGGCCACAGGACACTTCTT |
| **PLIN 2** | CCTCCTGTCCAACATCCAAG | GCATTGCGGAACACTGAGTA |
| **PLIN 3** | CCACCAATGTGAAGGACCA | GAAGGAGTGGATGCTGGAAA |
| **PLIN 4** | AATGAGCAACTTCGGAGCAC | CACCGTGTGACTTTGGACAG |
| **PLIN 5** | CCACAAGCTGGGTTCTTCAC | TGATCCACCTGCCTCAGACT |

**Key resources table**

| REAGENT or RESSOURCE | SOURCE | IDENTIFIER |
|---|---|---|
| **Antibodies** | | |
| MICU2 | Abcam | ab101465 |
| MICU1 | Sigma | HPA037479 |
| MCU | Abcam | ab272488 |
| OPA1 | BD Biosciences | 612607 |
| MFN2 | Cell signaling | #11925 |
| DRP1 (D8H5) | Cell signaling | #5391 |
| P-DRP1 (S616) | Cell signaling | #3455 |
| NDUFS2 | Abcam | ab192022 |
| SDHA | Abcam | ab137040 |
| UQCRC2 | Abcam | ab14745 |
| MT-CO1 | Abcam | ab14705 |
| **Chemicals, peptides and recombinant proteins** | | |
| CyQuant | Thermo Fisher Scientific | #35006 |
| Mitotracker | Invitrogen/Molecular Probes | M7514 |
| mt-riG6m | Li et al. 2020 (https://doi.org/10.1016/j.ceca.2020.102165) | |
| miGer | Li et al. 2020 (https://doi.org/10.1016/j.ceca.2020.102165) | |
| Nucleospin RNA plus Kit | Macherey–Nagel | 740984-250 |
| PrimeScript RT Reagent Kit | Takara | RR037A |
| SYBR Green Master kit | Takara | RR420L |
| Hypoxyprobe kit | Hypoxyprobe | HP1-1000Kit |
| Fura-2-AM | Thermo Fisher Scientific | #F1221 |
| MitoSox Red | Thermo Fisher Scientific | #M36008 |
| BCA protein assay kit | Thermo Fisher Scientific | #23227 |
| FAOBlue | Funakoshi (Japan) | FDV-0033 |
| **Inhibitors** | | |
| 2-Deoxy-D-glucose | Sigma-Aldrich | D8375 |
| 2NBDG | Abcam | ab146200 |
| UK5099 | MedChemExpress | HY-15475 |
| Rotenone | TOCRIS | CAS 83-79-4 |
| Oligomycin | Abcam | ab141829 |
| Trimetazidine | Sigma-Aldrich | 653322 |
| Etomoxir | Sigma-Aldrich | E1905 |
| Telaglenastat | MedChemExpress | HY-12248 |
| V-9302 | MedChemExpress | HY-112683 |
| BAPTA | Sigma-Aldrich | A1076 |
| MITOTEMPO | Sigma-Aldrich | SML0737 |
| **Treatments** | | |
| 5-Fluorouracil | Fluorouracile Accord 50mg/ml | |
| Oxaliplatin | Oxaliplatine Accord 5mg/ml | |
| **Software** | | |
| Image Lab | Bio Rad | |

| | |
|---|---|
| GraphPad Prism 6 | |
| Kaluza 1.3 software | Beckman Coulter |
| T-scratch software | |
| Image 4.4 software | Perkin Elmer |
| VisualSonics VevoLAZR System | FUJIFILM |
| ImageJ/Fiji | |
| Metamorph® 7.7 | Molecular Devices |
| Imaris 8.0® | Bitplane |
| SoftMax Pro 4.5.6 | |
| R 4.3.0 | R-Cran |

**Figure S1**

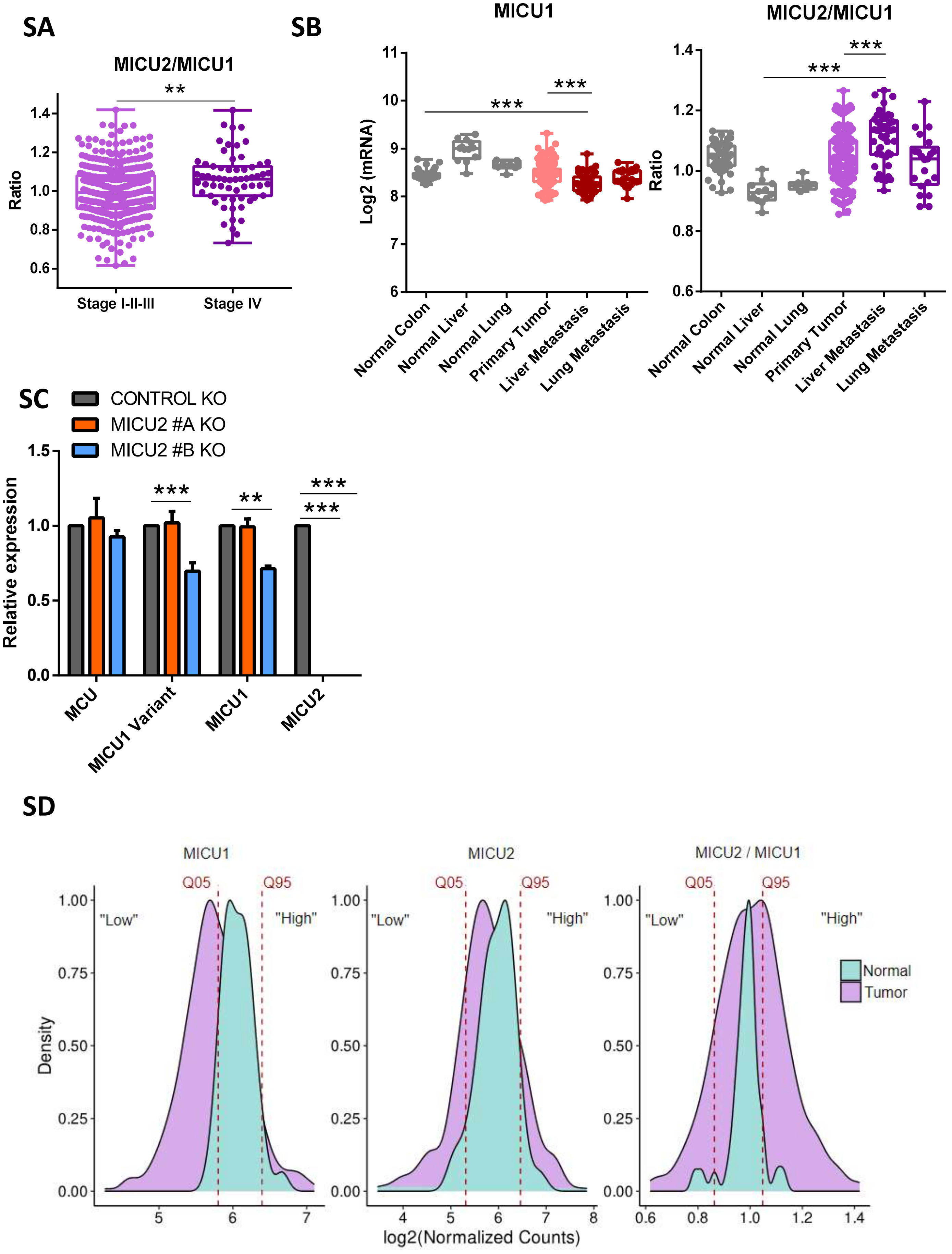
SA
MICU2/MICU1
**
Ratio
1.4
1.2
1.0
0.8
0.6
Stage I-II-III
Stage IV
SB
MICU1
Log2 (mRNA)
***
***
Normal Colon
Normal Liver
Normal Lung
Primary Tumor
Liver Metastasis
Lung Metastasis
MICU2/MICU1
Ratio
SC
CONTROL KO
MICU2 #A KO
MICU2 #B KO
Relative expression
MCU
MICU1 Variant
MICU1
MICU2
SD
MICU1
MICU2
MICU2 / MICU1
Q05
Q95
"Low"
"High"
Density
log2(Normalized Counts)
Normal
Tumor

**Figure S2**

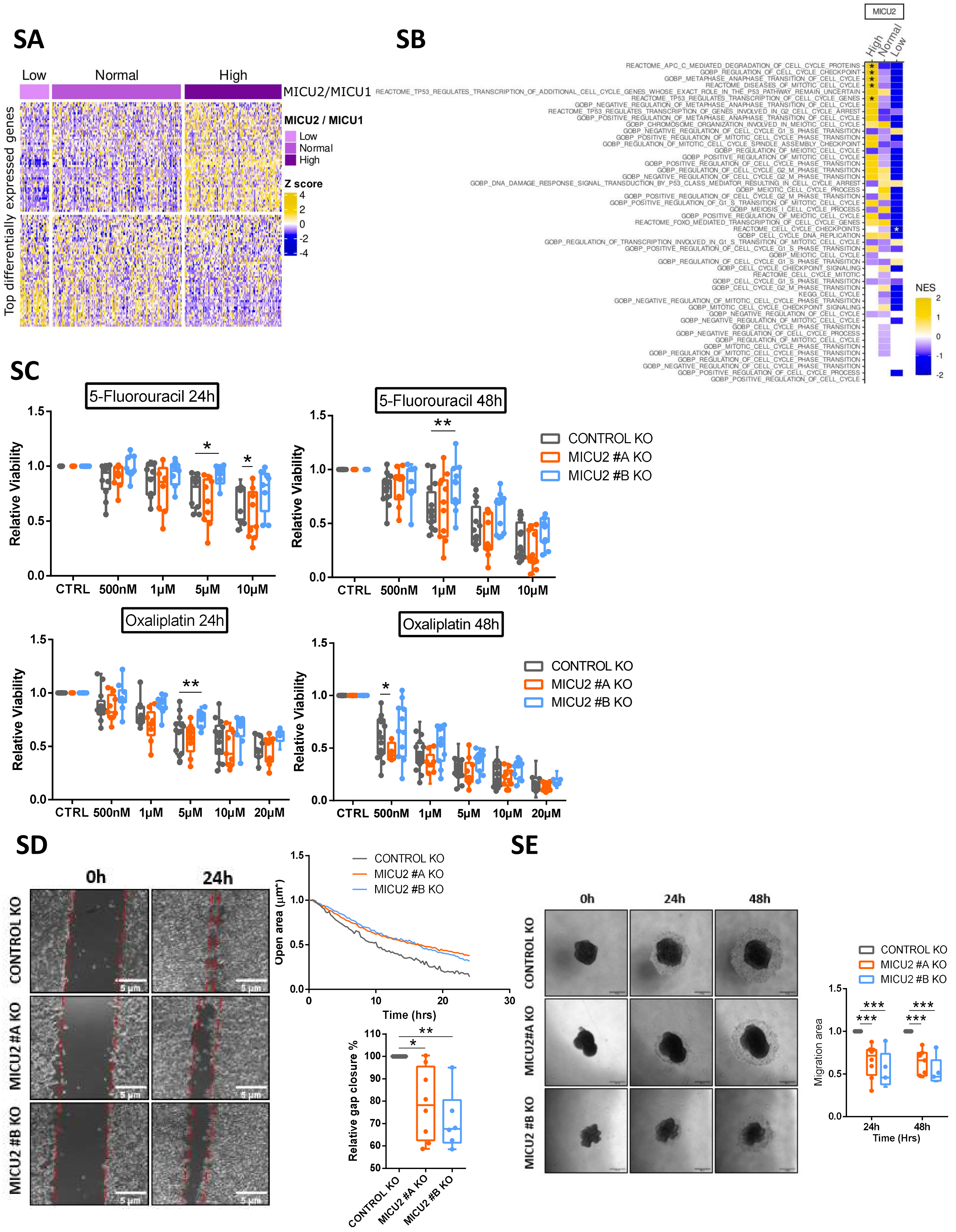
SA
Low
Normal
High
MICU2/MICU1
Top differentially expressed genes
MICU2 / MICU1
Low
Normal
High
Z score
4
2
0
-2
-4
SB
MICU2
High
Normal
Low
NES
2
1
0
-1
-2
SC
5-Fluorouracil 24h
5-Fluorouracil 48h
Oxaliplatin 24h
Oxaliplatin 48h
Relative Viability
CTRL
500nM
1μM
5μM
10μM
20μM
CONTROL KO
MICU2 #A KO
MICU2 #B KO
SD
0h
24h
CONTROL KO
MICU2 #A KO
MICU2 #B KO
Open area (μm²)
Time (hrs)
Relative gap closure %
SE
0h
24h
48h
CONTROL KO
MICU2#A KO
MICU2 #B KO
Migration area
Time (Hrs)

### Figure S3

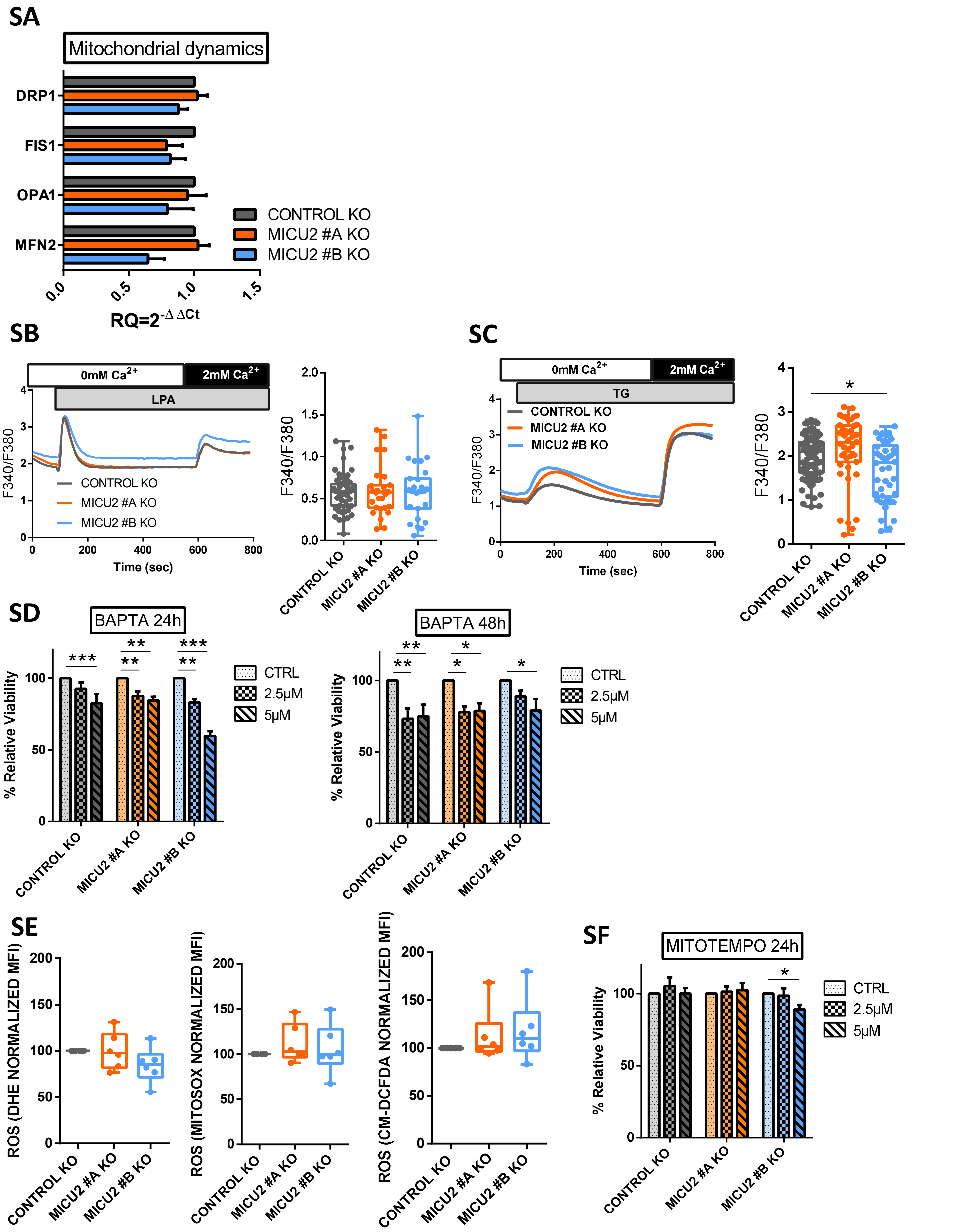
SA
Mitochondrial dynamics
DRP1
FIS1
OPA1
MFN2
CONTROL KO
MICU2 #A KO
MICU2 #B KO
0.0
0.5
1.0
1.5
RQ=2-ΔΔCt
SB
0mM Ca2+
2mM Ca2+
LPA
F340/F380
Time (sec)
CONTROL KO
MICU2 #A KO
MICU2 #B KO
SC
0mM Ca2+
2mM Ca2+
TG
CONTROL KO
MICU2 #A KO
MICU2 #B KO
F340/F380
Time (sec)
*
SD
BAPTA 24h
BAPTA 48h
% Relative Viability
CTRL
2.5μM
5μM
***
**
*
CONTROL KO
MICU2 #A KO
MICU2 #B KO
SE
ROS (DHE NORMALIZED MFI)
ROS (MITOSOX NORMALIZED MFI)
ROS (CM-DCFDA NORMALIZED MFI)
SF
MITOTEMPO 24h
% Relative Viability
CTRL
2.5μM
5μM

**Figure S4**

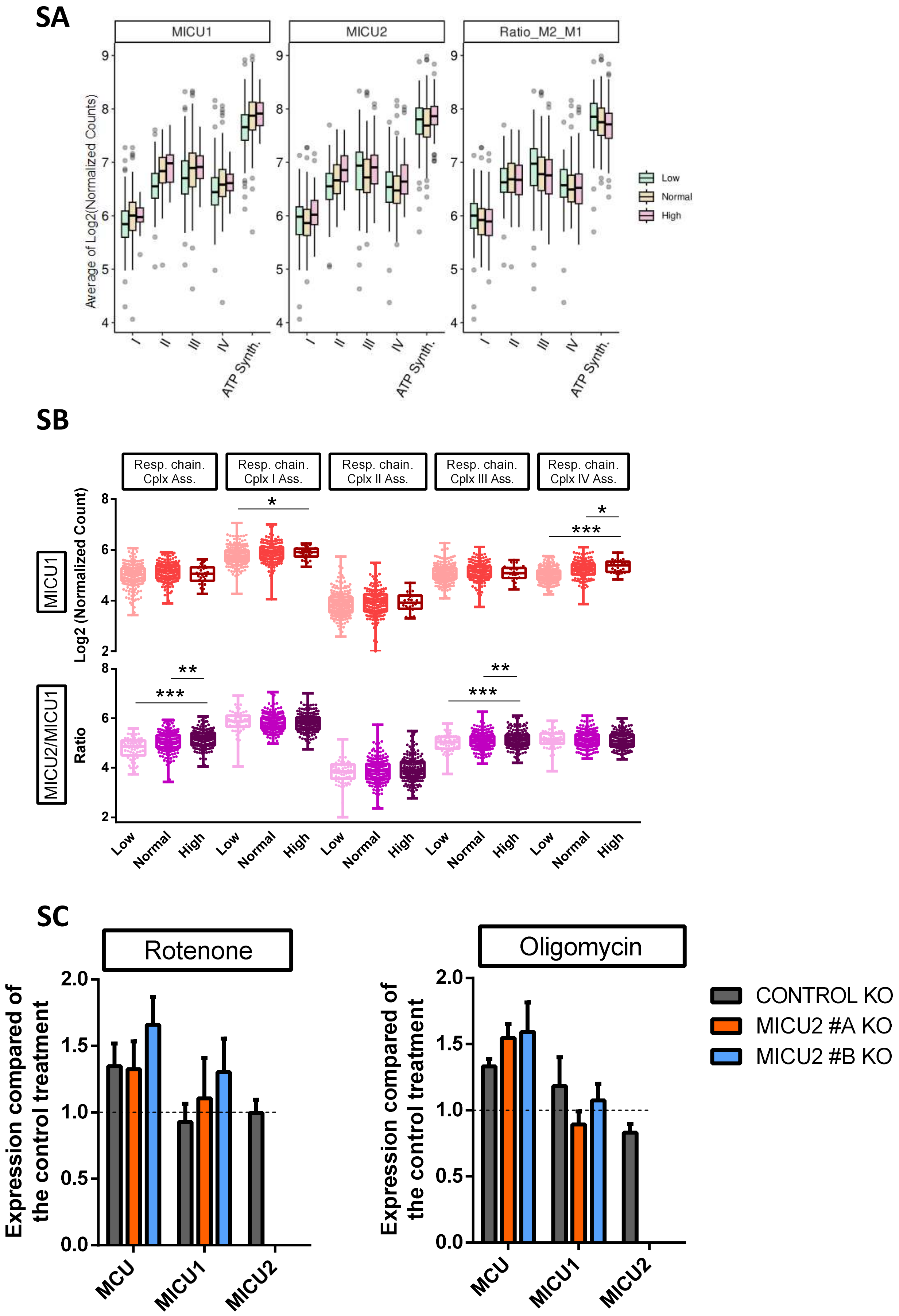
SA
MICU1
MICU2
Ratio_M2_M1
Average of Log2(Normalized Counts)
9
8
7
6
5
4
I
II
III
IV
ATP Synth.
Low
Normal
High
SB
Resp. chain. Cplx Ass.
Resp. chain. Cplx I Ass.
Resp. chain. Cplx II Ass.
Resp. chain. Cplx III Ass.
Resp. chain. Cplx IV Ass.
MICU1
Log2 (Normalized Count)
MICU2/MICU1
Ratio
8
6
4
2
*
***
**
Low
Normal
High
SC
Rotenone
Oligomycin
Expression compared of the control treatment
2.0
1.5
1.0
0.5
0.0
MCU
MICU1
MICU2
CONTROL KO
MICU2 #A KO
MICU2 #B KO

**Figure S5**

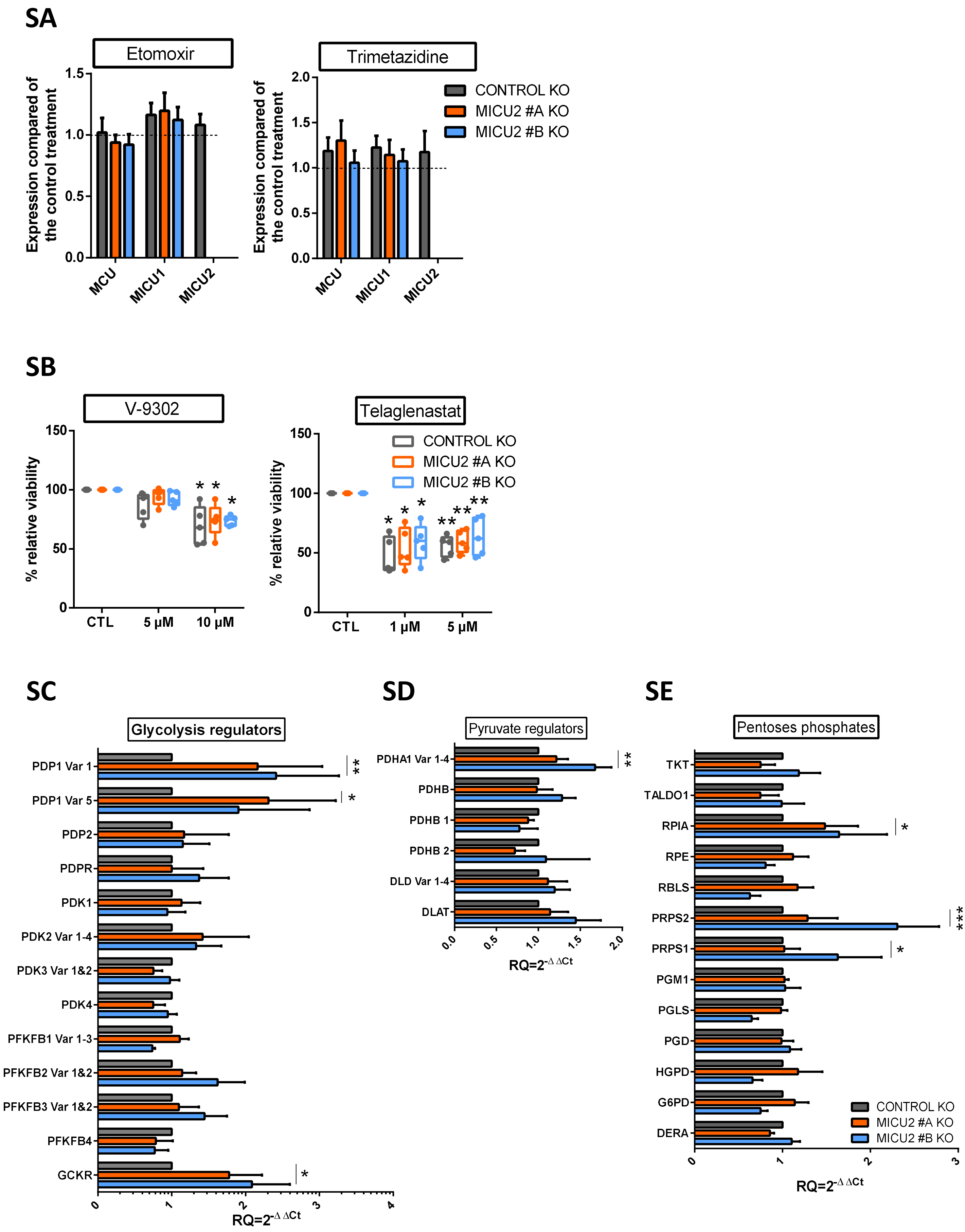
SA
Etomoxir
Expression compared of the control treatment
1.5
1.0
0.5
0.0
MCU
MICU1
MICU2
Trimetazidine
2.0
1.5
1.0
0.5
0.0
CONTROL KO
MICU2 #A KO
MICU2 #B KO
SB
V-9302
% relative viability
150
100
50
0
CTL
5 µM
10 µM
Telaglenastat
CTL
1 µM
5 µM
SC
Glycolysis regulators
PDP1 Var 1
PDP1 Var 5
PDP2
PDPR
PDK1
PDK2 Var 1-4
PDK3 Var 1&2
PDK4
PFKFB1 Var 1-3
PFKFB2 Var 1&2
PFKFB3 Var 1&2
PFKFB4
GCKR
RQ=2^-ΔΔCt
SD
Pyruvate regulators
PDHA1 Var 1-4
PDHB
PDHB 1
PDHB 2
DLD Var 1-4
DLAT
SE
Pentoses phosphates
TKT
TALDO1
RPIA
RPE
RBLS
PRPS2
PRPS1
PGM1
PGLS
PGD
HGPD
G6PD
DERA

**Figure S6**

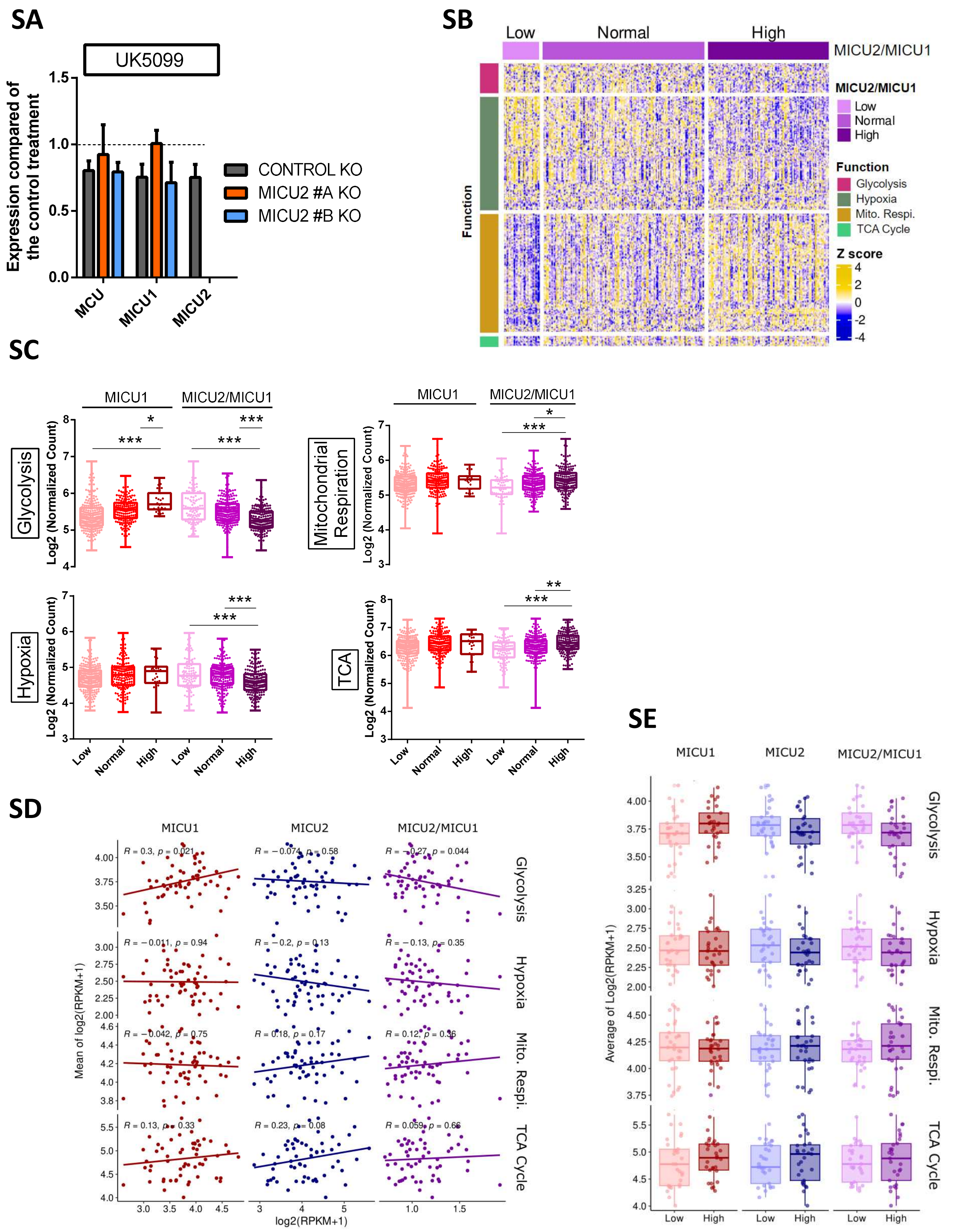

SA
UK5099
Expression compared of the control treatment
CONTROL KO
MICU2 #A KO
MICU2 #B KO
MCU
MICU1
MICU2
SB
Low
Normal
High
MICU2/MICU1
Function
Glycolysis
Hypoxia
Mito. Respi.
TCA Cycle
Z score
SC
Glycolysis
Mitochondrial Respiration
Hypoxia
TCA
Log2 (Normalized Count)
SD
MICU1
MICU2
Mean of log2(RPKM+1)
log2(RPKM+1)
R = 0.3, p = 0.021
R = −0.074, p = 0.58
R = −0.27, p = 0.044
R = −0.011, p = 0.94
R = −0.2, p = 0.13
R = −0.13, p = 0.35
R = −0.042, p = 0.75
R = 0.18, p = 0.17
R = 0.12, p = 0.36
R = 0.13, p = 0.33
R = 0.23, p = 0.08
R = 0.059, p = 0.66
SE
Average of Log2(RPKM+1)
